# A ROS-Ca^2+^ signalling pathway identified from a chemical screen for modifiers of sugar-activated circadian gene expression

**DOI:** 10.1101/2021.11.20.469363

**Authors:** Xiang Li, Dongjing Deng, Gizem Cataltepe, Ángela Román, Carolina Cassano Monte-Bello, Aleksandra Skyricz, Camila Caldana, Michael J Haydon

## Abstract

Sugars are essential metabolites for energy and anabolism that can also act as signals to regulate plant physiology and development. Experimental tools to disrupt major sugar signalling pathways are limited. We have performed a chemical screen for modifiers of activation of circadian gene expression by sugars to discover pharmacological tools to investigate and manipulate plant sugar signalling. Using a library of commercially available bioactive compounds, we identified 75 confident hits that modified the response of a circadian luciferase reporter to sucrose in dark-adapted seedlings. We validated the transcriptional effect on a subset of the hits and measured their effects on a range of sugar-dependent phenotypes for 13 of these chemicals. Chemicals were identified that appear to influence known and unknown sugar signalling pathways. Pentamidine isethionate (PI) was identified as a modifier of a sugar-activated Ca^2+^ signal that acts downstream of superoxide in a metabolic signalling pathway affecting circadian rhythms, primary metabolism and plant growth. Our data provide a resource of new experimental tools to manipulate plant sugar signalling and identify novel components of these pathways.

## Introduction

Cells depend on sugars to generate energy and to build the molecules required for cellular form and function. Sugars can also act as signalling molecules with various roles in regulating growth and development, physiological processes, metabolic feedback and modulating abiotic or biotic stress responses (Rolland *et al*., 2006). Plants generate their own sugars from photosynthesis. This dependence on light for energy supply creates specific challenges for plant cells, which must maintain these processes under both predictable and unpredictable fluctuations in the growth environment. This requires multiple sugar signalling pathways to coordinate dynamic supply and demand throughout the plant.

There are four well-recognised sugar signalling pathways in plants. HEXOKINASE 1 (HXK1) is responsible for the first enzymatic step in glycolysis but has glucose signalling functions independent of its enzymatic activity (Moore *et al*., 2003). G-protein signalling plays a role in extracellular glucose sensing and cell proliferation (Chen *et al*., 2003; Urano *et al*., 2012). TARGET OF RAPAMYCIN (TOR) kinase functions in numerous signalling pathways and is activated under C-replete conditions (Xiong *et al*., 2013). By contrast, Snf1 RELATED KINASE 1 (SnRK1) is active under C starvation (Baena-González *et al*., 2007). SnRK1 activity is inhibited by the signalling sugar trehalose-6-phosphate (T6P) (Zhang *et al*., 2009), which is very tightly connected to sucrose levels (Figueroa & Lunn, 2016). SnRK1, and perhaps also TOR, can directly affect activity of transcription factors (Xiong *et al*., 2013; Mair *et al*., 2015). HXK1 can localise to the nucleus and associate with DNA-binding complexes (Cho *et al*., 2006).

The critical importance of sugar signalling in plant cells makes genetic analysis of these pathways challenging. Loss-of-function mutants in *TOR* or *T6P SYNTHASE 1 (TPS1)* are embryo lethal (Eastmond *et al*., 2002; Menand *et al*., 2002) and a double mutant in both catalytic subunits of SnRK1 is not viable (Ramon *et al*., 2019). Therefore, most studies on these pathways have used hypomorphic mutants or inducible transgenic lines (Baena-González *et al*., 2007; Gómez *et al*., 2010; Xiong *et al*., 2013; Belda-Palazón *et al*., 2020). By contrast, growth effects in mutants in *HXK1* or *REGULATOR OF G-PROTEIN SIGNALLING 1 (RGS1)* are relatively minor, but both mutants are hyposensitive to growth inhibition by high exogenous sugar (Moore *et al*., 2003; Chen *et al*., 2006).

Although there is significant overlap between the cellular processes controlled by these sugar signalling pathways, particularly growth and energy metabolism, there are distinct features of their signalling outputs. For example, genetic experiments indicate additive effects of *hxk1-3* and *rgs1-2* mutants (Huang *et al*., 2015), suggesting functionally distinct pathways. TOR regulates proteostasis, autophagy and cell cycle control by sugars (Burkart & Brandizzi, 2021), whereas SnRK1 controls responses to energy deprivation and regulation of iron homeostasis (Peixoto *et al*., 2021).

The circadian clock is a gene regulatory network that integrates external and intrinsic signals to coordinate biological rhythms according to daily and seasonal changes in the environment. Photoautotrophic metabolism requires feedback between C availability and the circadian oscillator to optimise plant growth and fitness. Sugars affect circadian rhythms in Arabidopsis in several ways. C status contributes to entrainment, the process of setting the circadian clock (Haydon *et al*., 2013), and measurement of photoperiod (Liu *et al*., 2021). Reduced photosynthesis lengthens circadian period, which can be supressed by supplying sugar (Haydon *et al*., 2013). Period adjustment by sugars requires T6P-SnRK1 signalling affecting transcription of *PSEUDO RESPONSE REGULATOR 7 (PRR7)* (Frank *et al*., 2018). Sugars can also affect amplitude of specific oscillator components. One mechanism occurs by post-transcriptional control of GIGANTEA (GI) and requires F-box protein ZEITLUPE (ZTL) (Haydon *et al*., 2017).

Circadian rhythms rapidly dampen in seedlings released into continuous darkness without supplied sugar. Application of sucrose to dark-adapted seedlings can re-initiate circadian rhythms and the phase is set according to the time of sugar application (Dalchau *et al*., 2011). This transcriptional response to sugar does not require GI and the signalling processes are not known. This simple assay provides a sensitive technique to define sugar responses in the absence of light signals. Transcriptome analysis of this response revealed a role for superoxide, a reactive oxygen species (ROS), in promoting circadian gene expression and growth by sugar (Román *et al*., 2021). To further understand this transcriptional response to sugar, we screened the Library of Pharmacologically Active Compounds (LOPAC; Sigma) for chemicals that modify the response of a circadian reporter to sucrose in dark-adapted seedlings. From a list of 75 confident hit compounds, we selected 15 compounds to further characterise their effects on sugar-dependent processes. We identified two compounds that contribute to a sugar-activated ROS-Ca^2+^ signalling pathway that affects circadian rhythms, primary metabolism and plant growth. Our data provide a resource of pharmacological tools to manipulate sugar signalling in plants and has revealed opportunities to define new components of metabolic signalling.

## Materials and Methods

### Plant materials and growth conditions

Transgenic reporter lines in *Arabidopsis thaliana* for *COLD, CIRCADIAN RHYTHM AND RNA BINDING 2 (CCR2)* promoter:*LUCIFERASE (LUC)* (Doyle *et al*., 2002), *DARK INDUCIBLE 6 (DIN6)p:LUC* (Frank *et al*., 2018), *CaMV 35Sp:AEQUORIN (AEQ)* (Dalchau *et al*., 2010)*CIRCADIAN CLOCK ASSOCIATED 1 (CCA1)p*:*LUC* (CS9382), *TIMING OF CAB 1 (TOC1)p:LUC* (Nakamichi *et al*., 2005) are in Col-0. *35Sp:LUC* (CS9966) is in Ws-2. The *glucose insensitive 2 (gin2-1) abscisic acid 2 (aba2-1), constitutive triple response 1 (ctr1-12)* with a *CCA1p:LUC* transgene have been described (Haydon *et al*., 2013). *CCA1p:LUC* was introduced by crossing into *g protein alpha subunit 1(gpa1-4)* (Jones *et al*., 2003), *gtp binding protein 1(agb1-4)* (Ullah *et al*., 2003), and *rgs1-2* (Chen *et al*., 2003). The *g-protein gamma subunit 1(agg1-1) agg2-1 agg3-3* triple mutant (Thung *et al*., 2012) and *casein kinase a 1 (cka1-1) cka2-1 cka3-1* triple mutant (Wang *et al*., 2014).

For sterile culture, seeds were surface sterilised (30% (v/v) bleach, 0.02% (v/v) Triton X-100), washed three times in sterile water and sown on modified Hoagland media (HM) (Haydon *et al*., 2012) or half-strength Murashige and Skoog media (½ MS) (Sigma), solidified with 0.8% (w/v) agar Type M (Sigma). Seeds were chilled for 2 d at 4ºC and grown in 12 h light (80-100 µmol m^-2^ s^-1^):12 h dark at constant 20ºC (L:D).

### Chemical screen

Seven d old *CCR2p:LUC* seedlings grown in L:D on HM were wrapped in aluminium foil at dusk. Under dim green light, individual 10 d old seedlings were transferred in the afternoon to 96 well white LUMITRAC plates (Greiner) containing 250 µl HM with 0.1% DMSO or 25 µM LOPAC chemical (Sigma) and dosed with 1 mM D-luciferin, K^+^ salt (Cayman). Eighty LOPAC chemicals were included in each plate, plus positive and negative controls. Each plate was prepared in triplicate. After 84 h in darkness (subjective dawn), 25 µl of 10% (w/v) sucrose or mannitol and luciferase was measured at 1 h intervals for 24 h in the dark using orbital scan mode in a LUMIstar Omega plate reader fitted with a Microplate stacker (BMG Labtech). HiTSeekR (List *et al*., 2016)was used to identify compounds that significantly altered peak luminescence at 12 h after sucrose application after removing 47 data series from the total 4,224 deemed as false negatives. The raw data was log_2_ transformed and normalised with robust *z*-score method for general signal difference correction and inter-plate comparison before calculation of strictly standardised mean difference (SSMD) values.

### Luciferase assays

For sugar response assays, 7 d old seedlings grown in L:D on ½ MS were wrapped in aluminium foil at dusk. After 72 h, seedlings were transferred to ½ MS containing chemicals in 96 well LUMITRAC plates under dim green light and sugars were added 12 h later at subjective dawn. 1 mM D-luciferin was applied at least 12 h before commencing luminescence measurements using orbital scan mode in a LUMIstar Omega plate reader with Microplate stacker (BMG Labtech). Circadian rhythms were measured in 10 d old *CCA1p:LUC* and *TOC1p:LUC* seedlings grown on ½ MS in L:D and transferred at Zeitgeber Time 0 (ZT0) to 25-well imaging plates (Ting *et al*., 2022) containing media with DMSO or chemical before imaging luminescence with a Photon Counting System (HRPCS5, Photek) in continuous light (40 µmol m^-2^ s^-1^) or continuous low light (<10 µmol m^-2^ s^-1^) provided by red (640 nm) and blue (470 nm) LED lights.

### Quantitative RT-PCR

Total RNA was extracted from ∼30 mg snap frozen tissue with ISOLATE II RNA Plant Kit (Meridian Bioscience). cDNA was prepared from 0.5 µg DNase-treated RNA in 10 µl reactions of Tetro cDNA synthesis kit (Meridian Bioscience) using oligo(d)T primer. 10 µl PCR reactions were performed in technical duplicate with SensiFAST SYBR No-ROX (Meridian Bioscience) with 4 ng cDNA and 300 nM primers (Table S1) on a CFX96 Real-time PCR System (BioRad). Mean PCR reaction efficiencies were calculated for each primer pair with LinRegPCR (Ruijter *et al*., 2009) and used to calculate gene expression levels (PCR_efficiency^-Ct^).

### Growth assays

Germination was measured in non-chilled, surface sterilised Col-0 seeds sown onto ½ MS with DMSO or chemical and mannitol or sucrose at ZT0 and immediately placed in L:D. Radicle emergence was scored at 2 or 3 timepoints per day for 4 d. For hypocotyl and root length measurements, Col-0 was grown in L:D on ½ MS for 2 d and transferred to media containing DMSO or chemical with 30 mM mannitol or sucrose at ZT0, wrapped in aluminium foil and grown on vertical plates for 5 d. Hypocotyl and root lengths were measured from photographs with ImageJ (NIH).

### Pigment quantification

Chlorophyll was extracted from 12 7 d old seedlings in 250 µl methanol and quantified by absorbance spectrophotometry (Porra, 1989). Anthocyanin was extracted from 5 9 d old seedlings in 250 µl methanol:1% (v/v) HCl and quantified by absorbance spectrophotometry (Chen *et al*., 2019).

### ROS measurements

L-012 luminescence and nitroblue tetrazolium (NBT) staining were performed in dark-adapted seedlings in liquid ½ MS as described (Román *et al*., 2021).

### Aequorin experiments

For chemical response assays, ∼10 8 d old *35Sp:AEQ* seedlings grown in L:D on ½ MS were transferred before dusk to 100 µl 5 µM coelenterazine h (Cayman) in 96 well LUMITRAC plates (Greiner). Luminescence was measured at 1 s intervals in the dark from subjective dawn using orbital scan mode in a LUMIstar Omega plate reader (BMG Labtech) and 50 µl of DMSO or chemicals were applied with an injector at 30 s for final concentration of 10 µM DPI or 25 µM PI. After 270 s, 150 µl discharge solution (1 M CaCl_2_, 10 % (v/v) ethanol) was injected and cytosolic Ca^2+^ concentration was calculated (Fricker *et al*., 1999). For sugar response assays, ∼10 7 d old *35Sp:AEQ* seedlings grown in L:D were transferred to 100 µl liquid ½ MS with 20 µM coelenterazine at dusk in 96 well LUMITRAC plates (Greiner) and wrapped in aluminium foil. After 84 h (subjective dawn), 50 µl ½ MS containing DMSO or chemicals were added for a final concentration of 5 µM DPI or 25 µM PI and luminescence was measured at 2 min intervals by orbital scan mode in a LUMIstar Omega plate reader (BMG Labtech). After 1 h, 75 µl sucrose or mannitol was added for a final concentration of 30 mM with DMSO, 5 µM DPI or 25 µM PI and luminescence was measured at 2 min intervals for 6 h.

### RNA-Seq

Fourteen d old Col-0 seedlings grown on ½ MS in L:D were transferred to media containing DMSO, 10 µM DPI, 25 µM PI, 12.5 µM AEG3482 or 2 µM Tyrphostin AG879 at ZT24, before lights on. Untreated control seedlings were collected at time of transfer and treated seedlings were collected at ZT2 and snap frozen in liquid N. Total RNA was extracted from biological triplicates with ISOLATE II RNA Plant kit (Meridian Bioscience). and quantified and qualified by Agilent 2100 Bioanalyzer, NanoDrop (ThermoFisher) and 1% agarose gel. 1 μg total RNA with RIN value above 7 was used for library preparation by Genewiz using NEBNext® UltraTM RNA Library Prep Kit for Illumina® and NEBNext Poly(A) mRNA Magnetic Isolation Module (NEB). Size selection of Adaptor-ligated DNA was then performed using AxyPrep Mag PCR Clean-up (Axygen), and fragments of ∼360 bp (with the approximate insert size of 300 bp) were recovered. Each sample was then amplified by PCR for 11 cycles using P5 and P7 primers. The PCR products were cleaned up using AxyPrep Mag PCR Clean-up (Axygen), validated using an Agilent 2100 Bioanalyzer, and quantified by Qubit 2.0 Fluorometer (Invitrogen). Libraries with different indices were multiplexed and loaded on an Illumina HiSeq instrument and sequenced using a 2×150bp paired-end (PE) configuration; image analysis and base calling were conducted by the HiSeq Control Software (HCS) + OLB + GAPipeline-1.6 (Illumina) on the HiSeq instrument. All libraries >20 M raw reads with Q30 and Q20 >95% and >98%, respectively.

SOAPnuke v2.1.0 (Chen *et al*., 2018) was used to remove adapter sequences and low-quality reads from the sequencing data. To identify all the transcripts, we used HiSAT2 v2.2.1 (Kim *et al*., 2015) to align the sequencing reads to the *Arabidopsis thaliana* Col-0 genome (TAIR10) and RSEM v1.2.22 (Li & Dewey, 2011) to quantify the gene expression level of each replicate. The differential expression analysis was performed by NOIseq v2.37.0 (Tarazona *et al*., 2015). DEGs were defined as log_2_ change > 0.6 and DE probability > 80%.

### Metabolomics

Fourteen d old Col-0 seedlings grown hydroponically were transferred to liquid ½ MS containing DMSO, 10 µM DPI, 25 µM PI, 12.5 µM AEG3482 or 2 µM Tyrphostin AG879 at ZT23. Tissue was collected for five biological replicates for untreated seedlings at ZT23 and treated seedlings at ZT24 (lights off), ZT1.5, ZT4, ZT8, ZT10.5, ZT12 (lights on), ZT22.5 and ZT24 (lights off) and snap frozen in liquid N. Fifty mg of ground tissue was used for MTBE:methanol:water 3:1:1 (v/v/v) extraction (Giavalisco *et al*., 2011). The 150 μl of the organic phase was dried and derivatized (Roessner *et al*., 2001). One μl of the derivatized samples were analyzed on a Combi-PAL autosampler (Agilent Technologies) coupled to an Agilent 7890 gas chromatograph coupled to a Leco Pegasus 2 time-of-flight mass spectrometer (LECO) (Weckwerth *et al*., 2004). Chromatograms were exported from Leco ChromaTOF software (version 3.25) to R software. Peak detection, retention time alignment, and library matching were performed using Target Search R-package (Cuadros-Inostroza *et al*., 2009).

Metabolites were quantified by the peak intensity of a selective mass. Metabolite intensities were normalized by dividing the fresh-weight, followed by the sum of total ion count as described previously (Huege *et al*., 2011; Giavalisco *et al*., 2011). Principal component analysis was performed using pcaMethods bioconductor package (Stacklies *et al*., 2007). The significances of metabolites were tested by *t*-test.

## Results

We exploited the re-initiation of circadian gene expression by sugar in dark-adapted seedlings to evaluate transcriptional sugar responses in Arabidopsis. Using a luciferase reporter system, this provides a sensitive, high-throughput assay to measure plant sugar responses in the absence of light. Topical application of 30 mM sucrose to dark-adapted seedlings is sufficient for a robust response of a circadian luciferase reporter *CCR2p:LUC* (Fig. 1a) and generates similar sugar concentrations to seedlings grown in the light (Fig. 1b). Sucrose is the most abundant and mobile soluble sugar in plant tissues, although similar reporter activity is also elicited by glucose, fructose and maltose (Fig. 1c).

**Figure 1.**
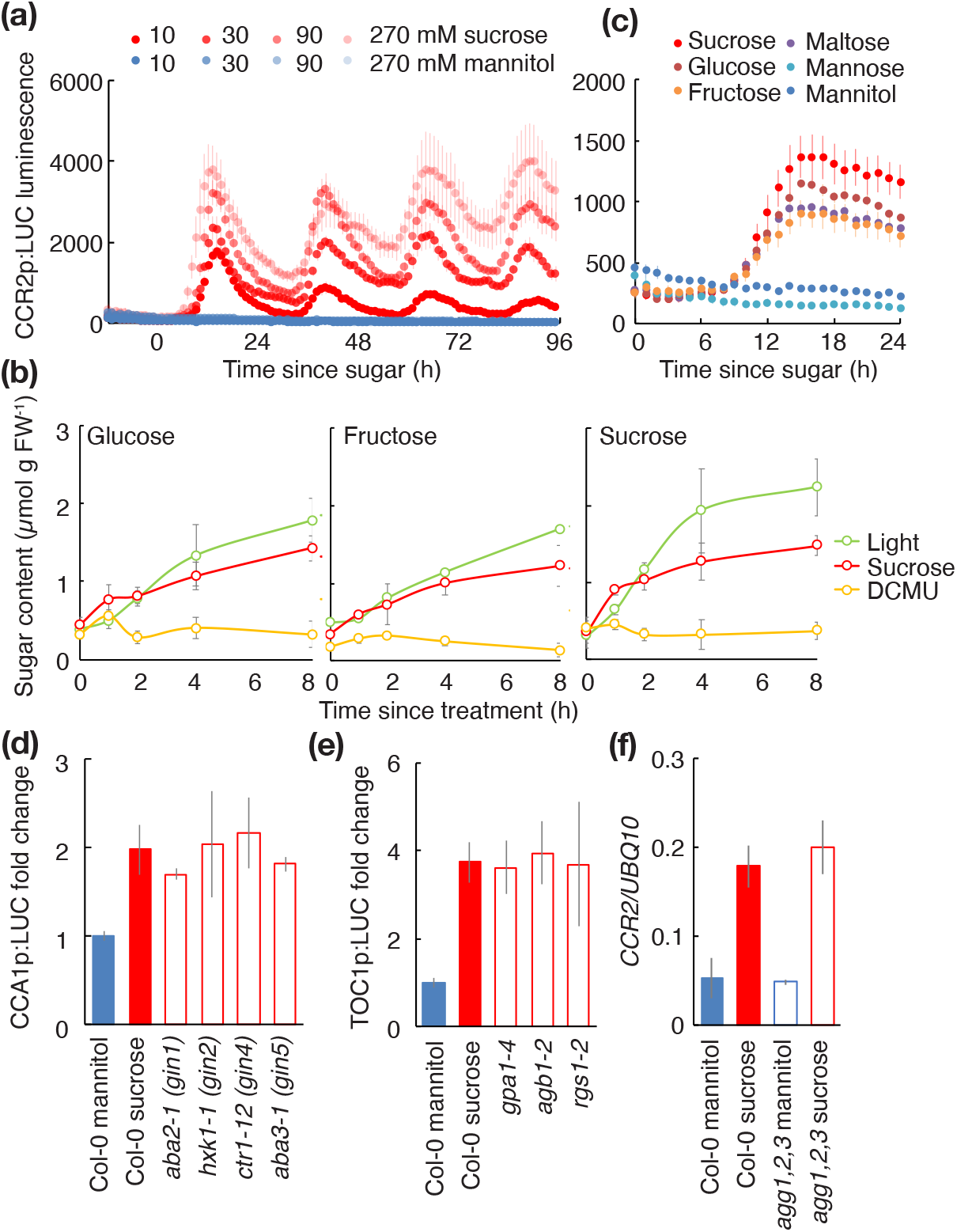
A sugar response assay in Arabidopsis seedlings. (a) Luciferase luminescence in dark-adapted *CCR2p:LUC* seedlings treated with indicated concentration of sucrose or mannitol (means ± SEM, n = 6). (b) Sugar content in dark-adapted seedlings treated with 30 mM sucrose or transferred into the light with or without DCMU (means ± SD, n = 4). (c) Luciferase luminescence in dark-adapted *CCR2p:LUC* seedlings treated with 30 mM sugars (means ± SEM, n = 8). (d) and (e) Fold change in luciferase reporter luminescence in dark-adapted wild-type (Col-0) or mutant seedlings treated with mannitol (blue) or sucrose (red) (means ± SD, n = 4). (f) *CCR2* transcript level, normalised to *UBQ10*, in dark-adapted wild-type (Col-0) or *agg1 agg2 agg3* mutant seedlings 12 h after treatment with mannitol (blue) or sucrose (red) (means ± SD, n = 3).

Numerous sugar-insensitive mutants have been identified in Arabidopsis that are resistant to growth inhibition by high concentrations of sugars in the media. We tested the transcriptional responses to sucrose in a collection of these mutants using our assay and observed similar responses to wild-type seedlings (Fig. 1d-f). Notably, these included mutants in *HXK1* and G-protein signalling, suggesting these sugar signalling pathways are not required for sugars to reinitiate circadian rhythms in dark-adapted seedlings.

To discover new components involved in plant sugar signalling, we performed a high-throughput screen for chemicals that modify the transcriptional response to sucrose. Dark-adapted *CCR2p:LUC* transgenic seedlings were treated with sucrose in the presence of 25 µM of each of 1280 chemicals from LOPAC (Sigma). This collection of commercially available chemicals have defined targets in mammalian cells, and mostly target signalling proteins such as receptors, kinases and ion channels. We used HiTSeekR (List *et al*., 2016) to calculate strictly standardised mean difference (SSMD) values. A cut-off of ±1 identified 146 chemicals that significantly modified the peak reporter activity (Table S2). To generate a higher confidence list of candidates, we used a stricter SSMD cut-off of ±1.28 which provided a list of 65 inhibitors and 10 enhancers of the reporter response to sucrose (Table 1).

**Table 1.**
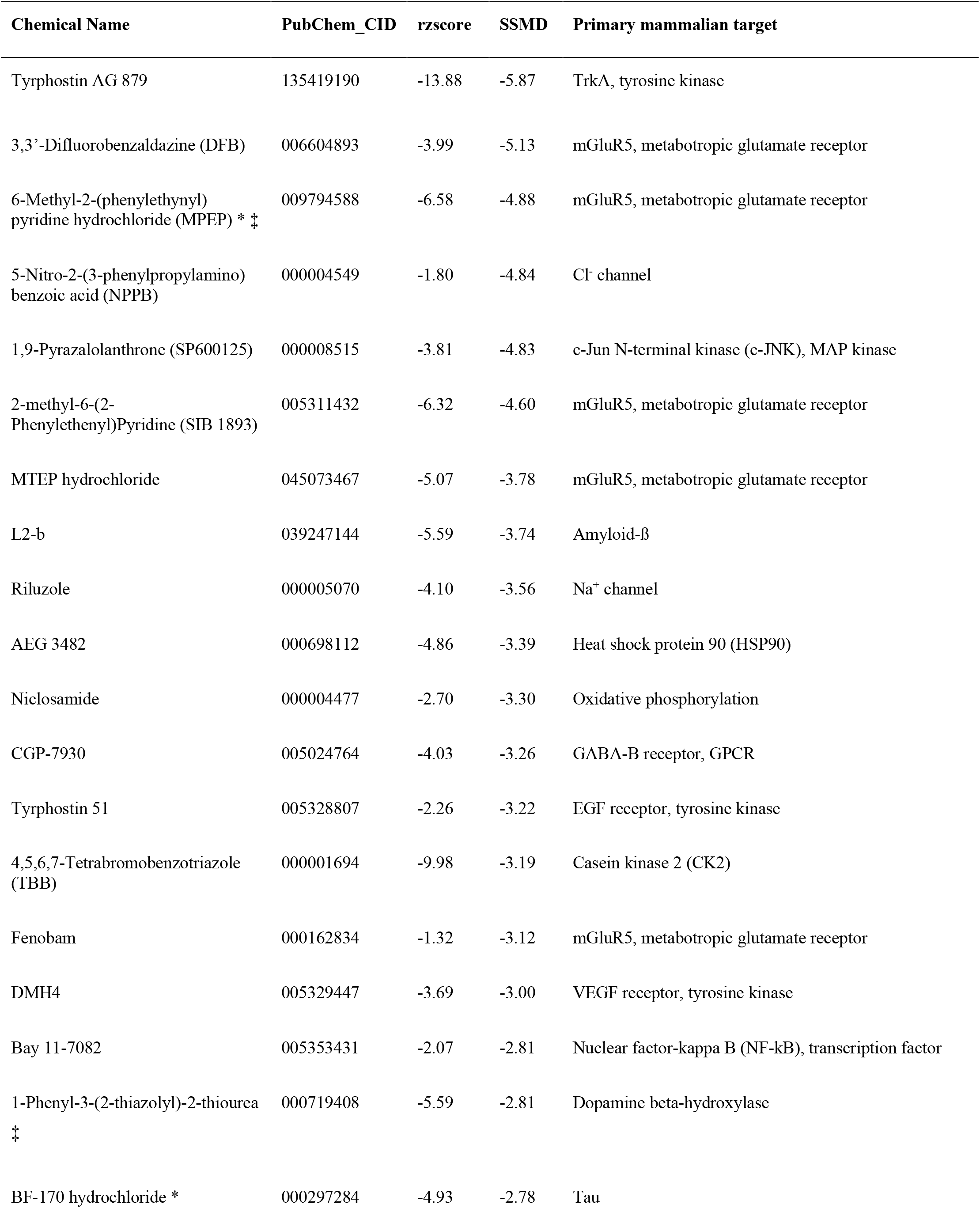
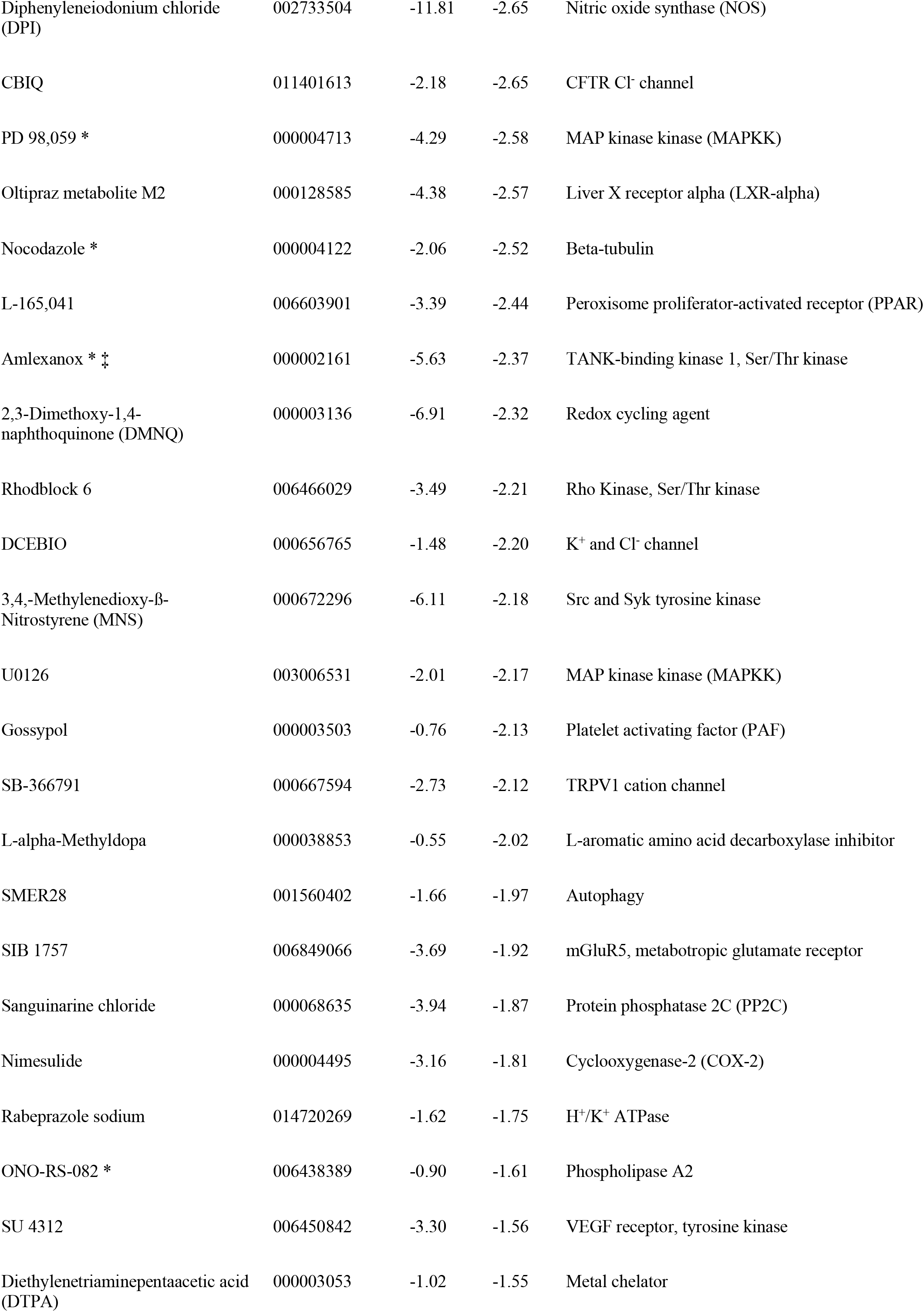
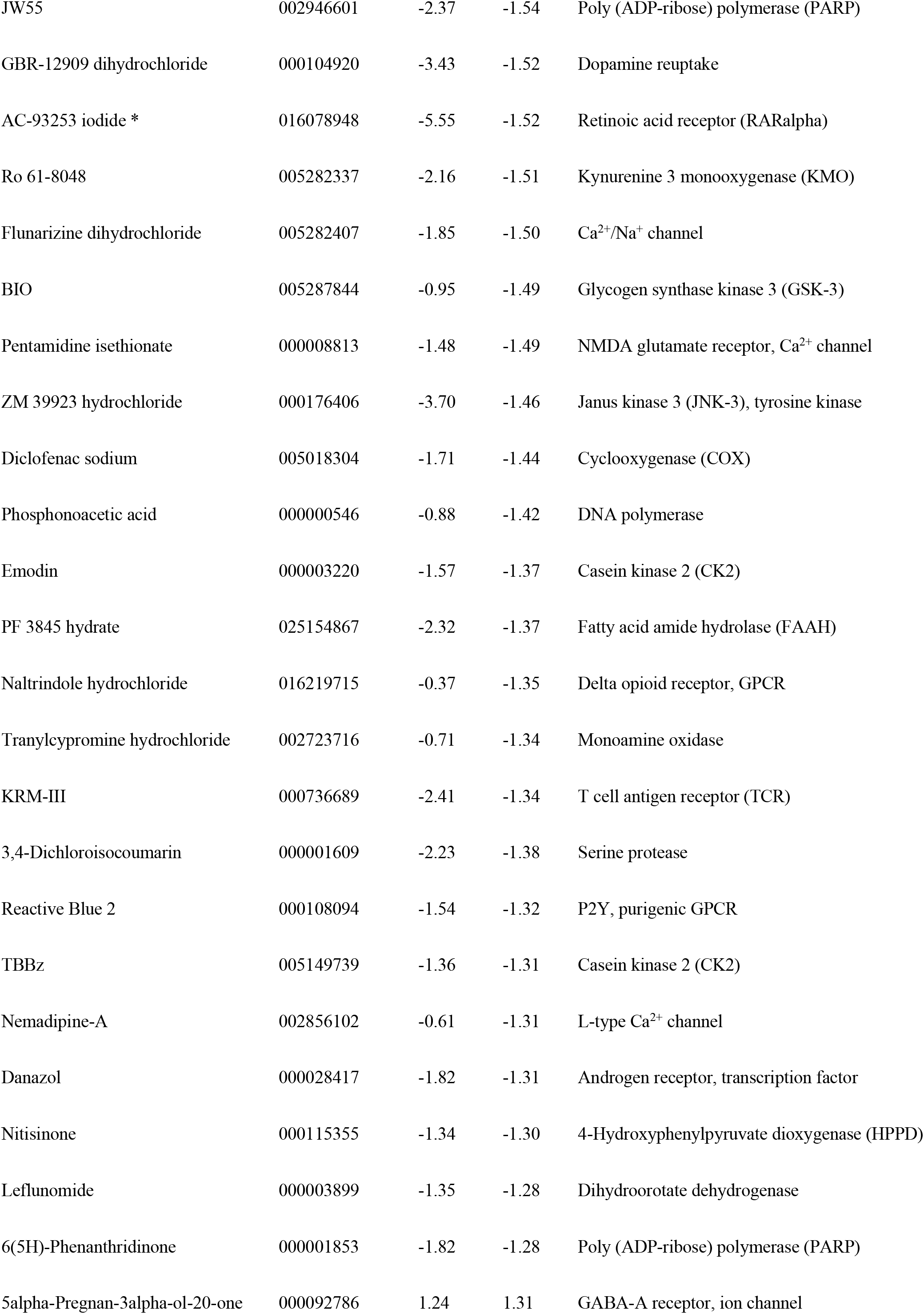
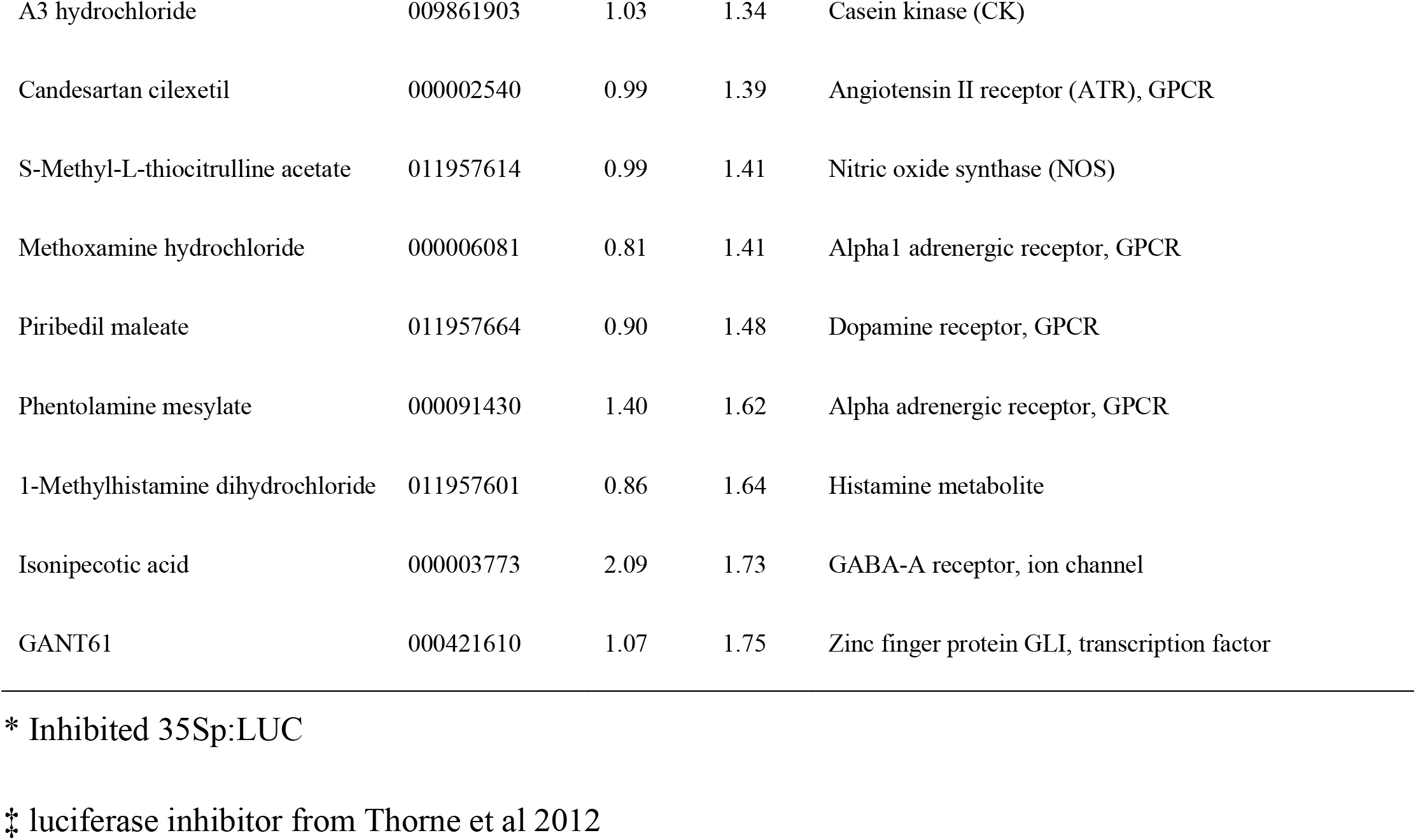
Chemical modifiers of *CCR2p:LUC* response to sucrose.

Greater than 10% of compounds present in chemical libraries are reported to exhibit inhibition of luciferase activity (Thorne *et al*., 2012). We re-screened 104 chemicals from LOPAC for their effect on luciferase activity at 25 µM using *35Sp:LUC* seedlings (Fig. S1). We identified two compounds that significantly enhanced luciferase luminescence and 11 compounds that significantly inhibited luminescence, indicating these might be false-positives identified in the primary screen.

We selected 15 chemicals (Fig. S2) for further analyses including diphenyleneiodinium (DPI), which we previously showed inhibits the transcriptional sugar response (Román *et al*., 2021), 6-methyl-2-(phenylethynyl)pyridine (MPEP), which strongly inhibited *35Sp:LUC* luciferase (Fig. S1), and AZD8055, a specific inhibitor of TOR (Montané & Menand, 2013), which is not in LOPAC but has been reported to affect circadian sugar responses in Arabidopsis (Zhang *et al*., 2019). We generated dose response curves for these 15 chemicals based on inhibition of *CCR2p:LUC* luminescence in dark-adapted seedlings treated with sucrose to determine a minimum effective concentration at which >80% inhibition is achieved (Fig. 2, Fig. S3). Although the primary screen being performed with 25 µM, effective concentrations ranged from 2-50 µM for these chemicals. By contrast, the maximum inhibition of the luciferase reporter achieved for the TOR inhibitor AZD8055 was only ∼50%.

**Figure 2.**
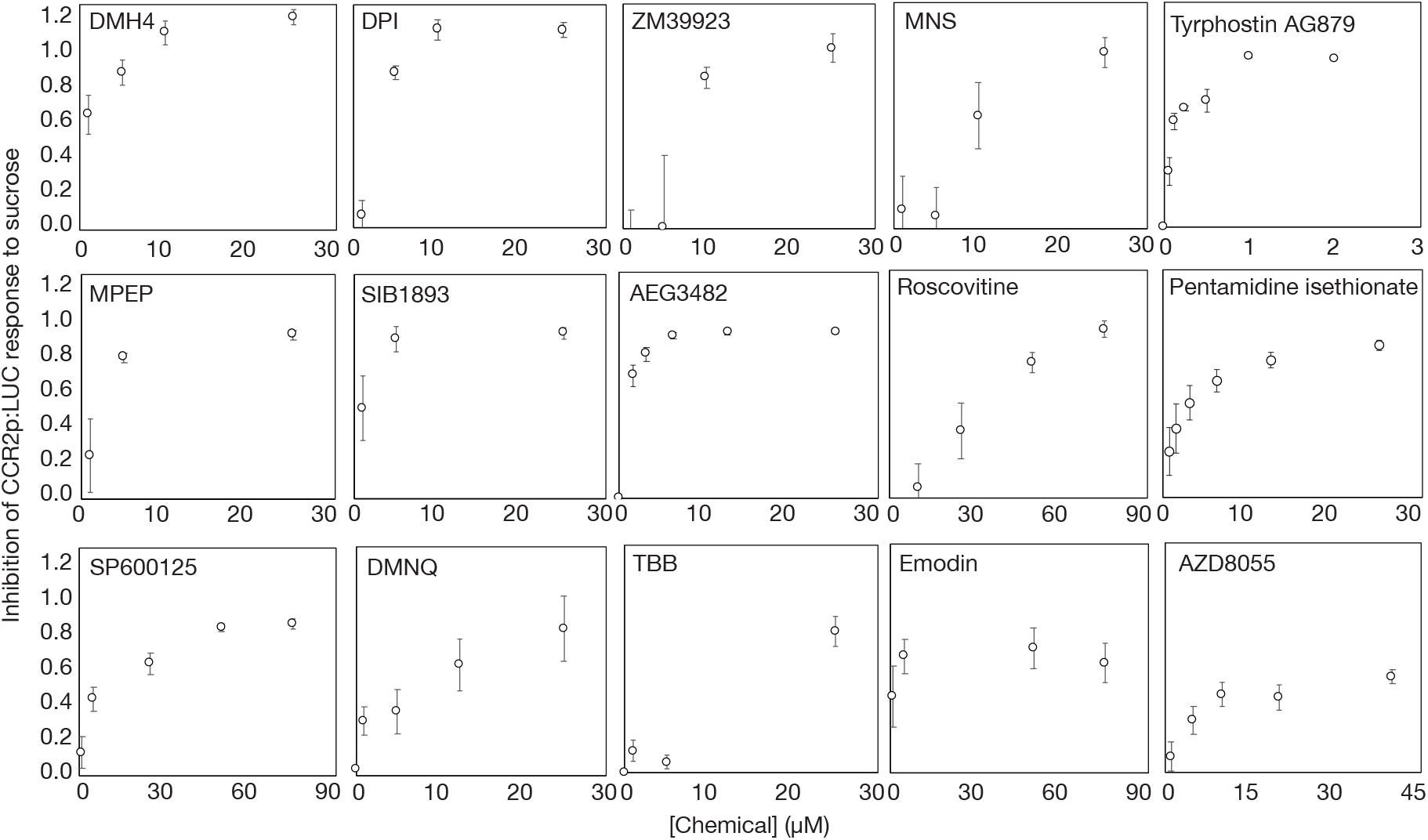
Dose response of transcriptional response to sucrose for 15 chemicals. Inhibition of peak luciferase luminescence in dark-adapted *CCR2p:LUC* seedlings after treatment with 30 mM sucrose in the presence of the indicated concentration of chemical (means ± SD, n = 6-12).

To validate the efficacy of these compounds on the transcriptional response to sugar, we again measured luciferase luminescence in *35Sp:LUC* seedlings at the minimum effective concentration determined from the dose curves (Fig. 3a) and measured *CCR2* transcript in dark-adapted seedlings treated with sucrose using qRT-PCR (Fig. 3b). This confirmed that all chemicals inhibited the upregulation of *CCR2* transcript by sucrose to some extent, except for two metabotropic glutamate receptor (mGluR5) inhibitors MPEP and SIB1893, which also significantly reduced luciferase luminescence in *35Sp:LUC* seedlings. We therefore excluded these two chemicals from further experiments. We also observed that 50 µM roscovitine increased 35Sp:LUC luminescence, despite inhibiting response of *CCR2p:LUC* and *CCR2* transcript to sucrose.

**Figure 3.**
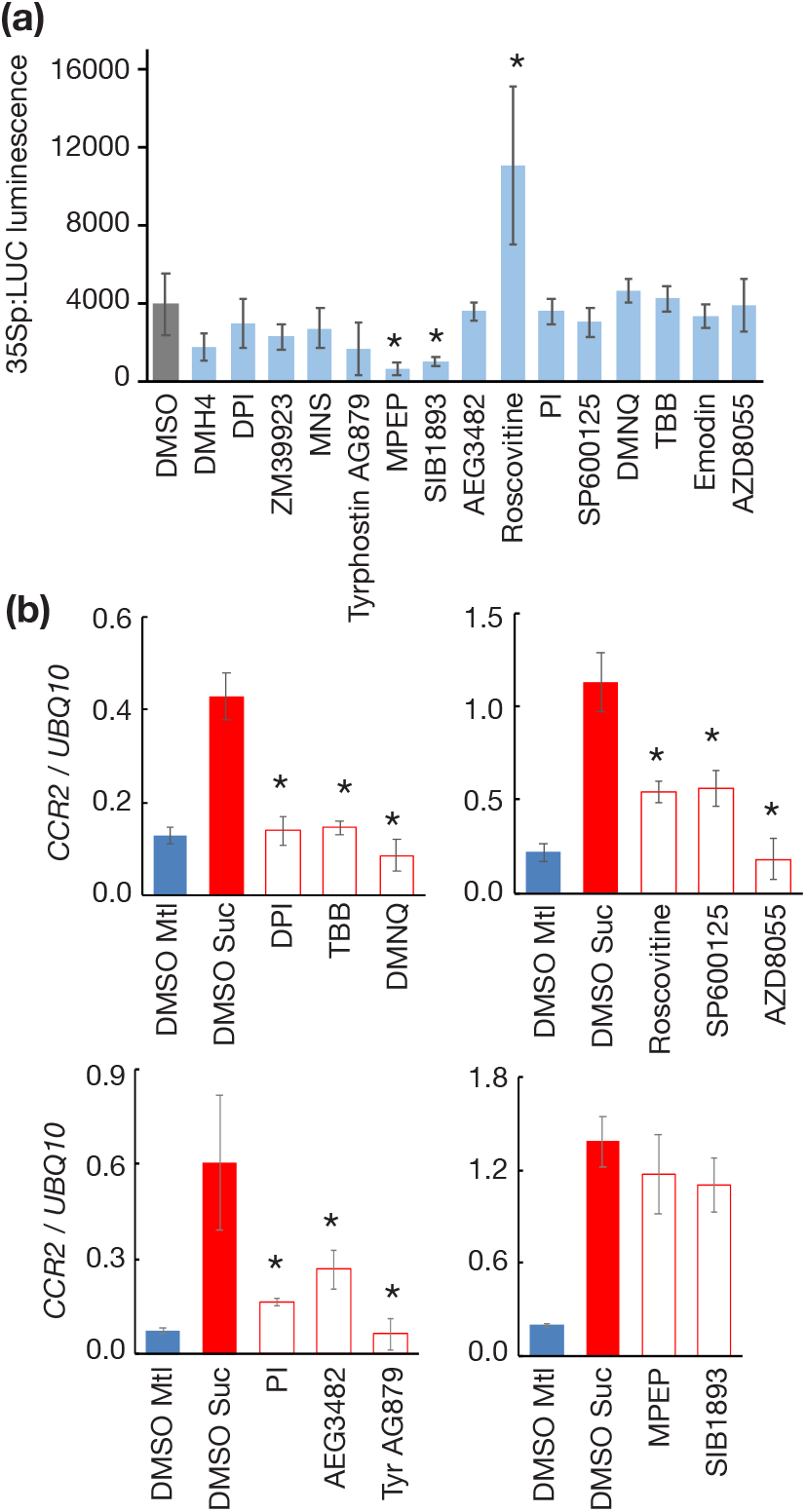
Validation of transcriptional effect of LOPAC chemicals. (a) Luciferase luminescence in *35Sp:LUC* seedlings, 16 h after transfer to media containing the minimum effective concentration of each chemical (means ± SD, n = 8; * p > 0.05, Bonferroni-corrected *t-*test) (b) *CCR2* transcript level, relative to *UBQ10*, in dark-adapted Col-0 seedlings 12 h after treatment with 30 mM mannitol (blue), 30 mM sucrose (red) or 30 mM sucrose in the presence of the minimum effective concentration of chemical (white) (means ± SD, n =4; * p > 0.05, Bonferroni-corrected *t-*test).

One of the validated chemicals, tetrabromobenzotriazole (TBB), is a mammalian casein kinase 2 inhibitor. CK2 is a regulator of the circadian clock and another CK2 inhibitor, dichlorobenzimidazole ribofuranoside (DRB), has been shown to lengthen circadian period in Arabidopsis (Portolés & Más, 2010). We tested whether DRB inhibits the response of *CCR2p:LUC* to sucrose in dark adapted seedlings and found no effect at the concentrations tested (Fig. S4a). Similarly, we could not detect a difference in *CCR2* transcript in a *cka1-1 cka2-1 cka3-1* mutants compared to wild type using qRT-PCR in the same assay (Fig. S4b). These data suggest that it is unlikely that TBB inhibits the transcriptional response to sucrose by targeting CK2 in Arabidopsis seedlings.

To build a picture of the broader influence of these selected chemicals on sugar-related processes, we tested their effects on a range of easily measurable phenotypes. Using the pre-determined minimum effective concentration for each chemical, we tested effects in the presence or absence of sucrose on germination of dormant seeds (Fig. S5), seedling biomass, chlorophyll content, hypocotyl length, root growth and anthocyanin content (Fig. S6). We observed a range of effects of each chemical on these phenotypes but detected similar patterns for multiple chemicals that might indicate common signalling pathways. To summarise these patterns, we plotted normalised effects of phenotypes on radar charts and ranked the chemicals by the sum of effects (Fig. 4a). Based on these radar charts, we noticed similar phenotypic patterns for DPI and pentamidine isethionate (PI), emodin and SP600125, and Tyrphostin AG879, DMH4 and ZM39923, which might indicate these chemicals affect proteins or signalling pathways with closely related functions.

**Figure 4.**
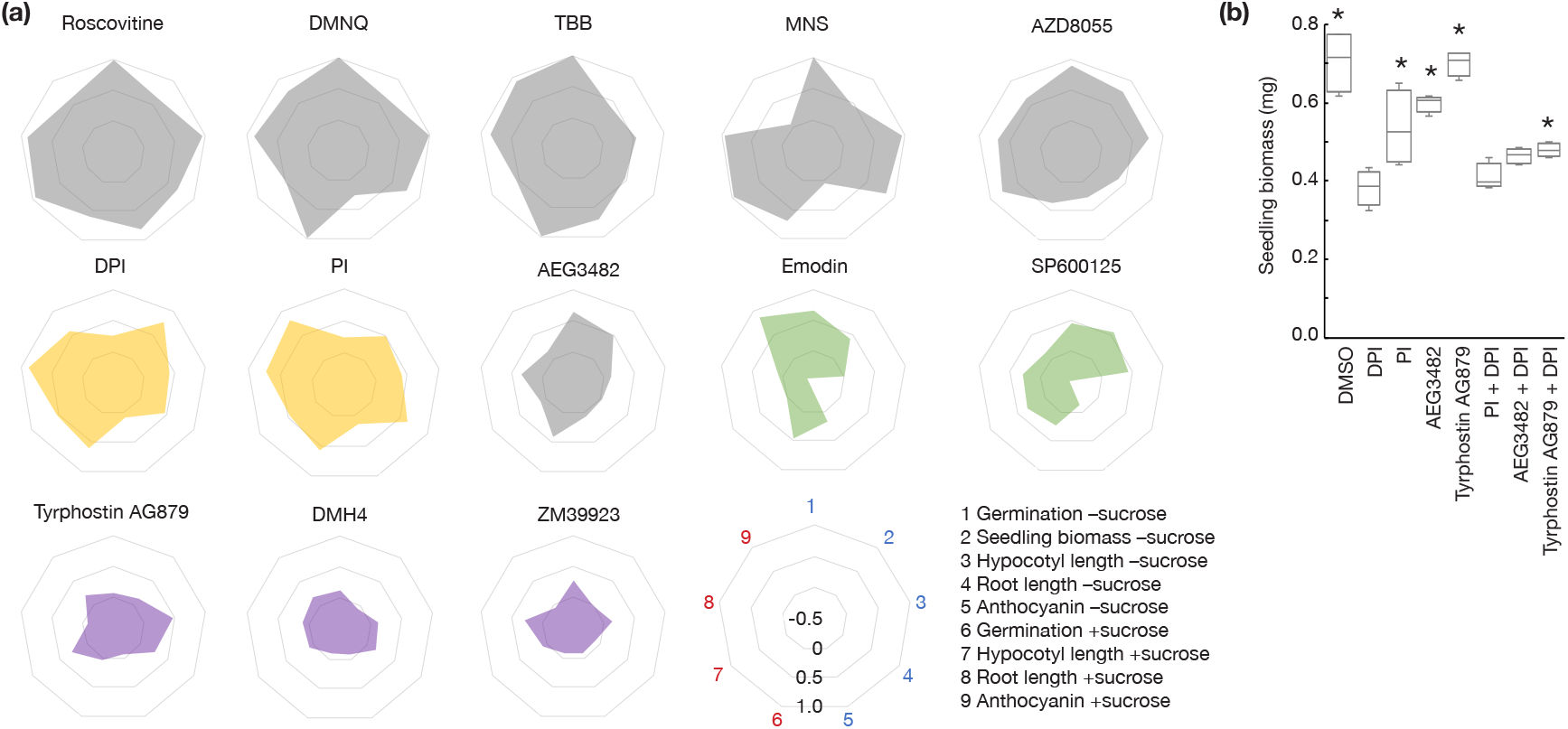
Summary of effects of LOPAC chemicals on sugar-related growth phenotypes. (a) Normalised effects of minimum effective concentration of 13 LOPAC chemicals on germination of dormant seeds on 30 mM mannitol (1) or sucrose (6), biomass of 7 d old seedlings on ½ MS (2), hypocotyl and root length of 7 d old dark-grown seedlings on 30 mM mannitol (3,4) or sucrose (7,8) and anthocyanin content in 9 d old seedlings grown for 2 d on 90 mM mannitol (5) or sucrose (9). Complete data are shown in Fig S4 and Fig S5. (b) Seedling biomass of 7 d old seedlings sown on ½ MS containing DMSO, 2 µM DPI, 5 µM PI, 2.5 µM AEG3482 or 0.5 µM Tyrphostin AG879 alone or in combination (n = 4 of 12 seedlings; * p < 0.05 compared to DPI, Bonferroni-corrected *t*-test).

We next tested phenotypic interactions between a selected number of chemicals. We hypothesised that chemicals that affect the same signalling pathway should be non-additive, whereas chemicals that affect distinct pathway might show a phenotypic interaction in combination. We germinated seeds on media containing chemicals alone or in combination at a 50% inhibitory concentration (Fig. 2) and measured biomass of 7 d old seedlings (Fig. 4b). DPI and PI in combination did not inhibit seedling growth compared to DPI alone, suggesting they affect the same pathway, consistent with the similar pattern for these chemicals in the radar charts. By contrast, Tyrphostin AG879 significantly suppressed the growth inhibition by DPI, indicating these chemicals might act on distinct pathways.

To measure the broader effects of these four chemicals on gene expression, we treated seedlings with each chemical at dawn and performed RNA-seq in seedlings after 2 h to capture short-term transcriptional effects at the beginning of the photoperiod. We identified 1899 differentially expressed genes (DEGs) between ZT0 and ZT2 (DMSO), but we detected between one and 16 DEGs between DMSO and any chemical (Table S2, Fig. S7a,b). This might be because the effect of inhibiting sugar signalling at dawn is diminished in the presence of abrupt light signalling. It’s also possible that since superoxide regulated genes are biased towards dusk (Román *et al*., 2021), the short-term effect of DPI is minimal around dawn. Nevertheless, more than two thirds of DEGs in chemical-treated seedlings verusus controls have been previously reported as sugar-regulated genes (Fig. S7b) (Xiong *et al*., 2013; Ganpudi *et al*., 2019; Román *et al*., 2021; Peixoto *et al*., 2021). *PATHOGEN AND CIRCADIAN CONTROLLED 1 (PCC1)*, which is upregulated after dawn, was identified as a DEG for DPI, PI and AEG3482, but not Tyrphostin AG879, providing further support that Tyrphostin AG879 acts on a distinct signalling pathway.

Half of the DEGs identified in Tyrphostin AG879-treated seedlings have been reported as DEGs in a *sesqiα2* SnRK1 hypomorphic mutant (Peixoto *et al*., 2021). We therefore tested whether Tyrphostin AG879 affects expression of a SnRK1 transcriptional marker, *DARK INDUCIBLE 6 (DIN6)*. Treatment of *DIN6p:LUC* seedlings at dusk with DPI, PI or AEG3482 either had no effect, or slightly inhibited luciferase luminescence, whereas Tyrphostin AG879 increased reporter activity (Fig. S7c). This suggests that Tyrphostin AG879 can activate SnRK1 activity, either directly or indirectly, which is consistent with the increased expression of SnRK1-regulated markers in Tyrphostin AG879-treated seedlings (Fig. S7b). An activation of the starvation response triggered by SnRK1 could explain how Tyrphostin AG879 counteracted the inhibition of seedlings growth by DPI (Fig. 4b).

To explore the effect of these four chemicals on primary metabolism, we performed metabolite profiling over a 24-h time-course following treatment with each chemical (Table S4). Principal component (PC) analysis of 63 metabolites over eight time-points revealed a similar trend for all four chemicals in the direction of change compared to DMSO-treated samples for PC1 and PC2, which together explained between 63-70% of variance (Fig. S8). The PC plot for Tyrphostin AG879 suggested a more pronounced effect on the primary metabolome around dusk, compared to the other three chemicals, and the effect of DPI appeared more pronounced than PI and AEG3482.

All four chemicals caused a significant increase in the levels of sucrose, fructose and glucose at more than one time-point. Sucrose was elevated around dawn and glucose and fructose elevated during the day (Fig 5a). The glycolysis intermediate glucose-6-phosphate was significantly lower in Tyrphostin AG879-treated seedlings at ZT10 and ZT12 and also reduced at ZT24 in seedlings treated with DPI and PI (Fig 5b). We observed significant increases in tricarboxylic acid (TCA) cycle intermediates, particularly citrate and malate, with the magnitude of the effect increasing with time for all four chemicals (Fig 5c). Branched-chain amino acids valine, leucine and isoleucine have been associated with HXK1 (Ganpudi *et al*., 2019) and TOR signalling (Cao *et al*., 2019) but we did not detect notable changes for any chemical (Fig. 5d). The difference between Tyrphostin AG879 and the other three chemicals detected around dusk in the PC analysis is likely explained by significantly reduced levels of amino acids at ZT10.5, particularly methionine, threonine and lysine (Fig. 5e), which are all synthesised from oxaloacetate in the TCA cycle.

**Figure 5.**
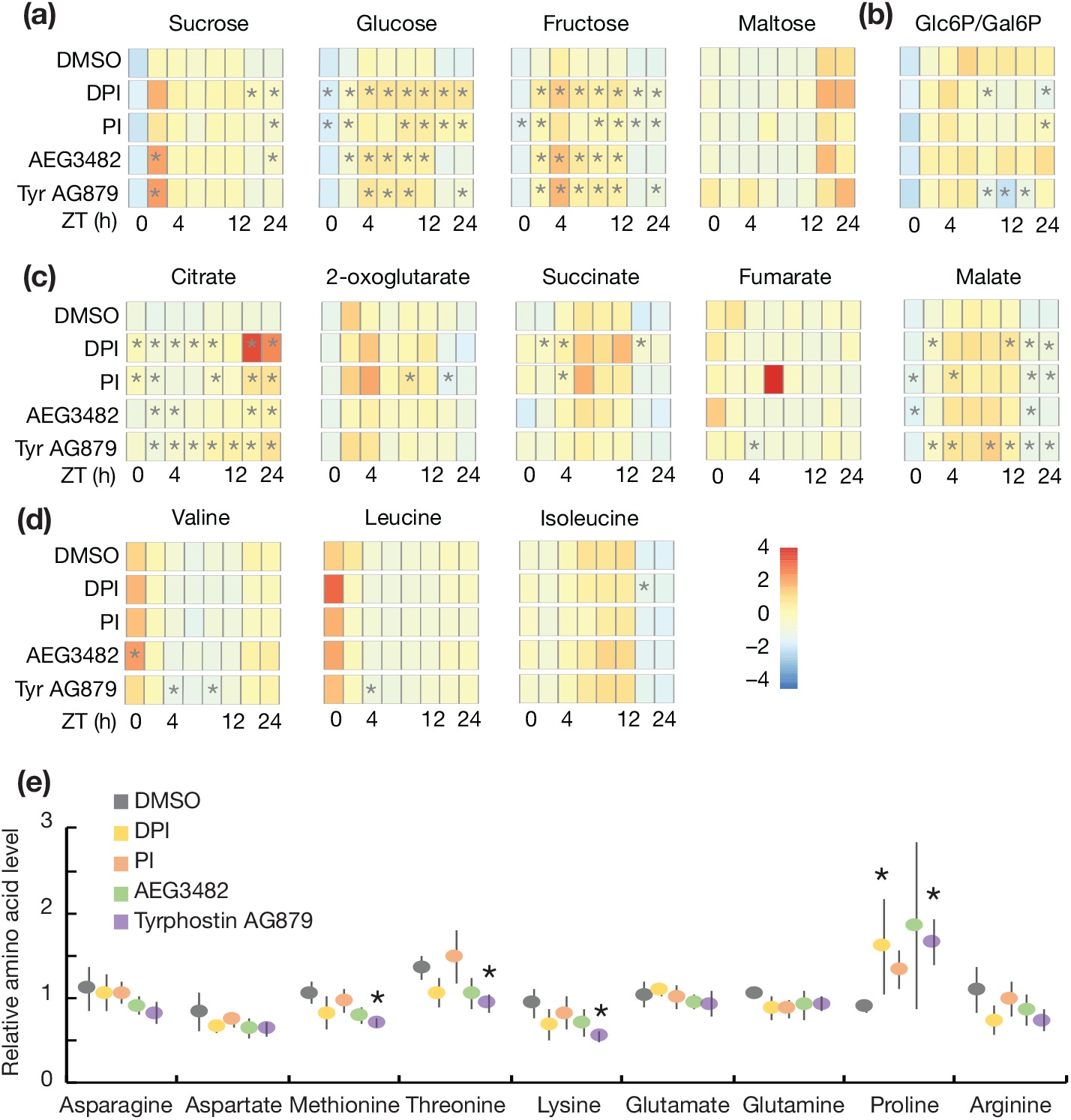
Primary metabolite levels in chemical-treated seedlings. (a-d) Heatmaps of relative metabolite levels in seedlings treated with DMSO or chemicals at ZT23 and sampled at ZT0, ZT1.5, ZT4, ZT8, ZT10.5, ZT12, ZT22.5 and ZT24 (n = 5; * p < 0.05 compared to DMSO, *t*-test). (e) Relative amino acid levels in DMSO- or chemical-treated seedlings at ZT10.5 (mean ± SD, n = 5; * p < 0.01 compared to DMSO, *t*-test).

We have previously shown that DPI inhibits accumulation of superoxide in dark-adapted seedlings treated with sucrose and affects expression of circadian gene expression (Román et al 2021). The similar phenotypic patterns between DPI and PI, suggest they might affect the same sugar signalling pathway. We measured the effect of DPI and PI on circadian rhythms using luciferase reporters and observed lengthening of circadian period by both chemicals (Fig. 6a). This effect is consistent with expectations for inhibition of sugar signalling into the clock, similar to effects of inhibiting sugar production from photosynthesis (Haydon *et al*., 2013). However, neither chemical inhibited the shortening of circadian period by sucrose (Fig. 6b), which operates by a SnRK1-dependent mechanism (Frank *et al*., 2018).

**Figure 6.**
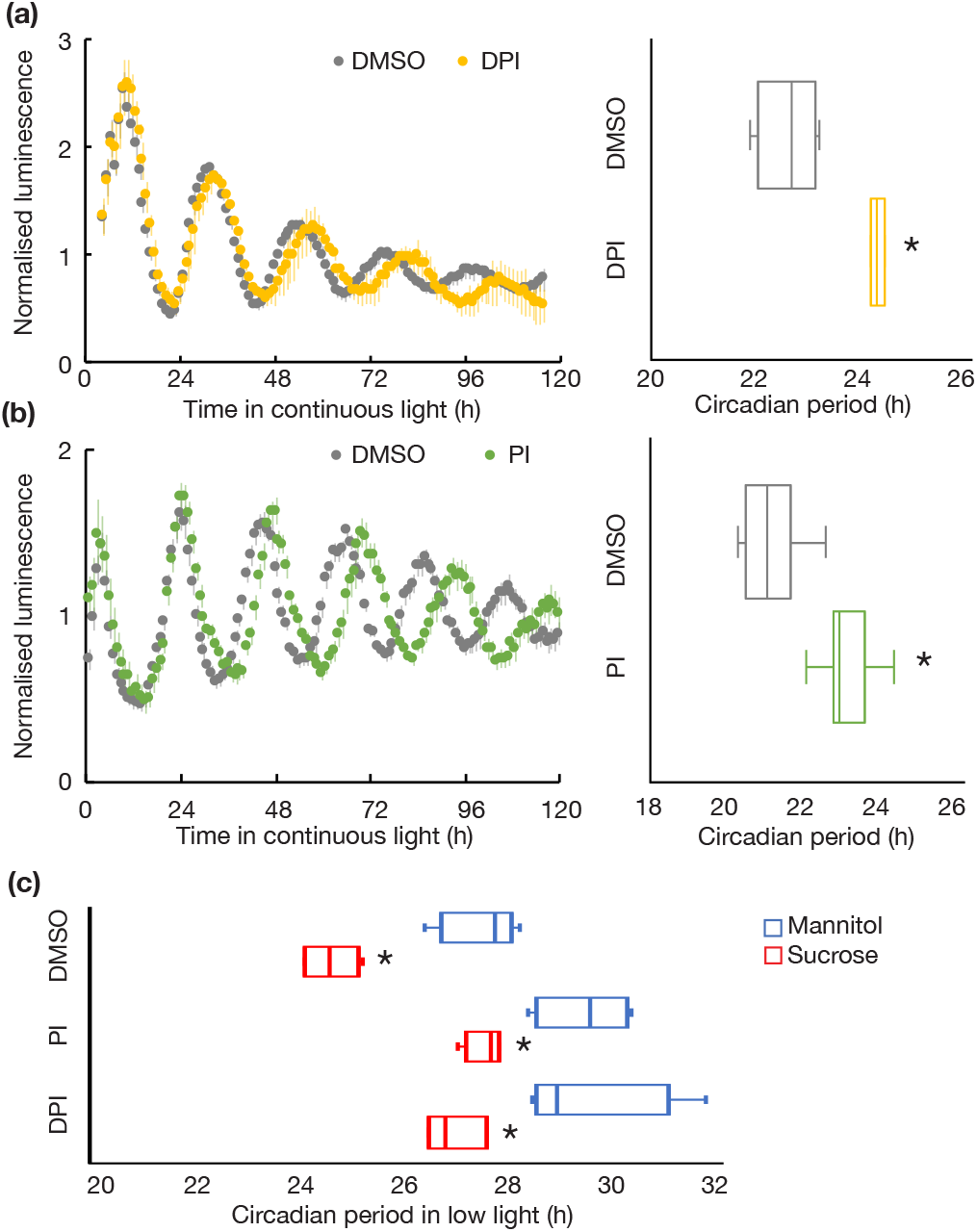
Effect of PI and DPI on circadian period. Normalised luciferase luminescence and circadian period estimates in (a) *TOC1p:LUC* (n = 4) or (b) *CCA1p:LUC* seedlings (n = 16) treated with DMSO, 10 µM DPI or 25 µM PI before transfer to continuous light (n = 4; * p < 0.05, Student’s *t*-test). (c) Circadian period estimates of *CCA1p:LUC* seedlings treated with DMSO, 10 µM DPI or 25 µM PI with 30 mM mannitol or sucrose before transfer into continuous low light (n = 4; * p < 0.05 compared to mannitol, Bonferroni-corrected *t*-test).

PI is an antagonist of N-methyl-D-aspartate (NMDA) glutamate receptors in mammalian cells, so we wondered if PI similarly inhibits Ca^2+^ signalling in Arabidopsis. We measured the effect of PI on *35Sp:AEQ*, a luminescent reporter for cytosolic Ca^2+^ concentration and observed a rapid shift in internal Ca^2+^ in Arabidopsis seedlings (Fig. 7a). By contrast, DPI did not affect 35Sp:AEQ in these experiments suggesting it does not have an immediate influence on cytosolic Ca^2+^ in these conditions.

**Figure 7.**
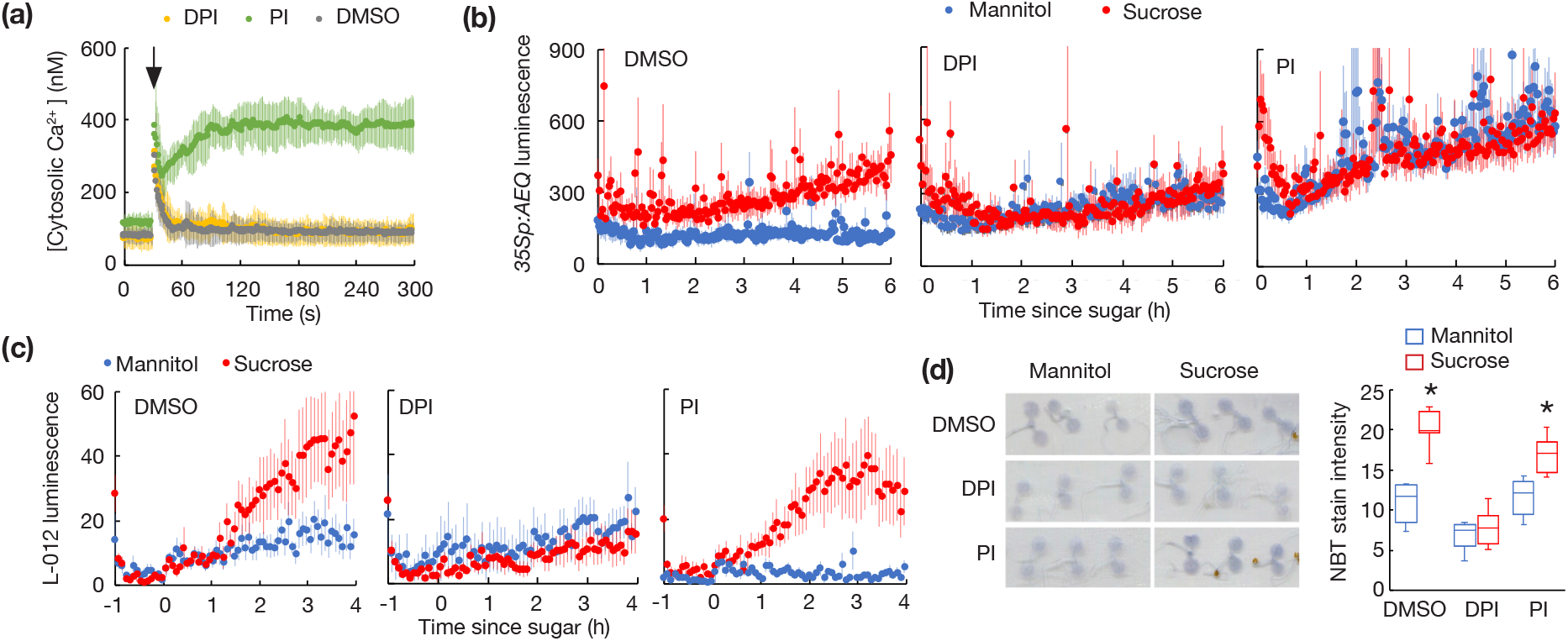
Effect of PI on elevation of cytosolic Ca^2+^ by sucrose. (a) Cytosolic Ca^2+^ concentration in *35Sp:AEQ* seedlings treated (arrow) with DMSO (control), 10 µM DPI or 25 µM PI (means ± SD, n = 6). (b) Aequorin luminescence in dark-adapted *35Sp:AEQ* seedlings treated with 30 mM sucrose or mannitol in the presence of DMSO, 5 µM DPI or 25 µM PI (mean ± SD, n = 6). (c) L-012 luminescence in dark-adapted Col-0 seedlings treated with 30 mM mannitol or sucrose in the presence of DMSO, 10 µM DPI or 25 µM PI (means ± SEM, n = 6). (d) Images and quantification of nitroblue tetrazolium (NBT) stain for superoxide in dark-adapted Col-0 seedlings 4 h after treatment with 30 mM mannitol or sucrose in the presence of DMSO, 5 µM DPI or 25 µM PI (n = 8; * p < 0.05 compared to mannitol, Bonferroni-corrected *t*-test).

Since PI strongly affects cytosolic Ca^2+^ concentration and inhibits transcriptional response to sucrose, we tested whether sucrose could induce a change in cytosolic Ca^2+^ concentration in dark-adapted seedlings. Application of sucrose to dark-adapted seedlings elevated 35Sp:AEQ luminescence compared to mannitol (Fig. 7b). This response was slower than elevation of superoxide observed in sucrose-treated seedlings (Román *et al*., 2021). Elevation of cytosolic Ca^2+^ concentration by sucrose was attenutated in DPI-treated seedlings. Similarly, although 35Sp:LUC luminescence was higher in PI-treated seedlings, there was no difference between sucrose and mannitol (Fig. 7b). This is consistent with the targets of these chemicals acting in the same sugar-activated Ca^2+^-dependent signalling pathway and suggests that the target of DPI acts upstream of the target of PI.

To test whether PI inhibits elevation of superoxide by sucrose, we measured L-012 luminescence and performed nitroblue tetrazolium (NBT) stains in dark-adapted seedlings. DPI strongly inhibited L-012 luminescence in sucrose-treated seedlings, but PI did not (Fig. 7c). Similarly, NBT stains indicated that, unlike DPI, PI did not prevent sucrose-activated superoxide accumulation (Fig. 7d). These data indicate that the target of PI acts downstream of the target of DPI. This supports the results of the 35Sp:AEQ experiments and suggests that PI inhibits a ROS-activated Ca^2+^-channel acting downstream of NADPH oxidases in sugar signalling pathway that regulates circadian gene expression.

## Discussion

We have used a chemical screen for modifiers of an effect of sucrose on the circadian clock to identify novel experimental tools to manipulate sugar signalling in Arabidopsis. From 75 hit molecules, we confirmed the effect of 13 chemicals and completed broad characterisation of the effects of these chemicals on key sugar-regulated processes including germination, growth and pigmentation. These experiments captured a broad picture of the patterns of phenotypic effects of these chemicals to identify potential shared relationships between them. We selected four chemicals for metabolite and transcriptome profiling, which suggested they affect at least two distinct pathways. Two compounds, DPI and PI, appear to act on a sucrose-activated ROS-Ca^2+^ signalling pathway, which might represent a novel metabolic signalling pathway in Arabidopsis affecting circadian gene expression and growth. Furthermore, this list of commercially available sugar signalling modifiers provides a resource that could be used to fill gaps in known and unknown metabolic signalling processes in plants.

The effect of Tyrphostin AG879 on the transcriptome suggested that it might be a modifier of SnRK1 or a SnRK1-related pathway, and we confirmed that this chemical could activate *DIN6p:LUC*, a reporter of SnRK1 activity. Although the number of DEGs was small in Tyrphostin AG879-treated seedlings around dawn, this is consistent with the small effect of *SnRK1α* -overexpression on the transcriptome and the apparently higher activity of SnRK1 at this time of day (Peixoto *et al*., 2021). A SnRK1 activator could be expected to inhibit the transcriptional response of *CCR2p:LUC* to sucrose in C-starved seedlings. Two other chemicals, DMH4 and ZM39923, had similar patterns of effects to Tyrphostin AG879, so it is possible that these also influence SnRK1-related signalling. Interestingly, all three chemicals are tyrosine kinase inhibitors.

The TOR kinase inhibitor, AZD8055, is not represented in LOPAC but we included it in our experiments because it was able to inhibit the transcriptional response to sucrose. Although the effect on the luciferase reporter was relatively weak, the effect on the transcript was strong. TOR is activated by sugar availability and influences circadian rhythms in Arabidopsis (Zhang *et al*., 2019), so it is possible other modifiers of plant TOR signalling could have been identified in our screen. Based on phenotypic effects, the most similar chemicals were roscovitine, a cyclin-dependent kinase inhibitor and TBB. Although TBB is a CK2 inhibitor, it appears this is not the mechanism by which is affects expression of *CCR2* in our experiments.

DPI and PI have very similar effects on the phenotypes we measured and their inhibition of growth is not additive, suggesting they act on the same pathway. DPI inhibits accumulation of superoxide in sucrose-treated seedlings, most likely by inhibiting NADPH oxidases at the plasma membrane (Román *et al*., 2021). PI is a NMDA receptor antagonist and therefore might target a Ca^2+^ channel in plant cells. Consistent with this, we found that PI induces a rapid change in cytosolic Ca^2+^ concentration. Application of sucrose to dark-adapted seedlings elevated cytosolic Ca^2+^, with a slightly slower response than the accumulation of superoxide. Since PI did not inhibit sucrose activated superoxide accumulation, these results suggest that the Ca^2+^ signal acts downstream of the superoxide signal and that PI targets a ROS-activated Ca^2+^channel. CYCLIC NUCLEOTIDE GATED Ca^2+^ CHANNELs (CNGCs) and ANNEXINs have been proposed to fulfil this role in plants (Demidchik, 2018). GLUTAMATE LIKE RECEPTORs (GLRs) have been shown to act upstream of NADPH oxidases (Kong *et al*., 2016). All these channels are members of large protein families, so genetic verification of any of these as the targets of PI will be challenging.

If our results suggest a novel sugar signalling pathway, a critical question is how the sugar is sensed. Since NADPH oxidases are present on the plasma membrane, this could be an extracellular sugar sensing mechanism. A role for RGS1 seems unlikely, since G-protein mutants are not affected in the sucrose response assay. Receptor-like kinases have been implicated in numerous extracellular signalling pathways, upstream of NADPH oxidases. This is a very large protein family, but many of these contain sugar-binding lectin domains (Sun *et al*., 2020). However, NADPH oxidases can be triggered by intracellular domains, so it’s also possible that sugars are sensed within cells. Alternatively, DPI might be inhibiting superoxide accumulation in a different location. It will be important to determine the cellular location of both the superoxide accumulation and Ca^2+^ signal in the future.

Confirming the direct targets of these chemicals will be important to define these signalling pathways. A forward genetic approach lacks sufficient specificity and the likelihood of isolating loss-of-function mutations in essential sugar signalling proteins will be small, since these are expected to be lethal. Reverse genetics is limited when there is functional redundancy within protein families. Therefore, chemical proteomics has the most promise (Hicks & Raikhel, 2014), although these techniques can be particularly challenging for membrane proteins. Nevertheless, this chemical screen has successfully identified a new set of potential tools to manipulate sugar signalling in plant cells. This provides opportunity to discover new signalling pathways in Arabidopsis and it will be informative to test the efficacy of these chemicals in other plant species.

## Acknowledgements

We thank Waheed Arshad and Heather Eastmond for technical assistance and Heather McFarlane and Antony Dodd for sharing seed. This research was funded by a Royal Society Research Grant (RG150144), the Botany Foundation and the University of Melbourne through the Research Grants Support Scheme to MJH and Melbourne Research Scholarships to XL and GC.

## Author Contribution

MJH conceived the study; all authors designed experiments and analysed data; XL, DD, GC, AR, CCMB and MJH performed experiments; XL and MJH wrote the manuscript; all authors edited and approved the manuscript.

## Data Availability

Raw sequencing files and processed data for RNA-seq have been deposited in the NCBI Gene Expression Omnibus [GSE188596].

---

**Figure S1.**
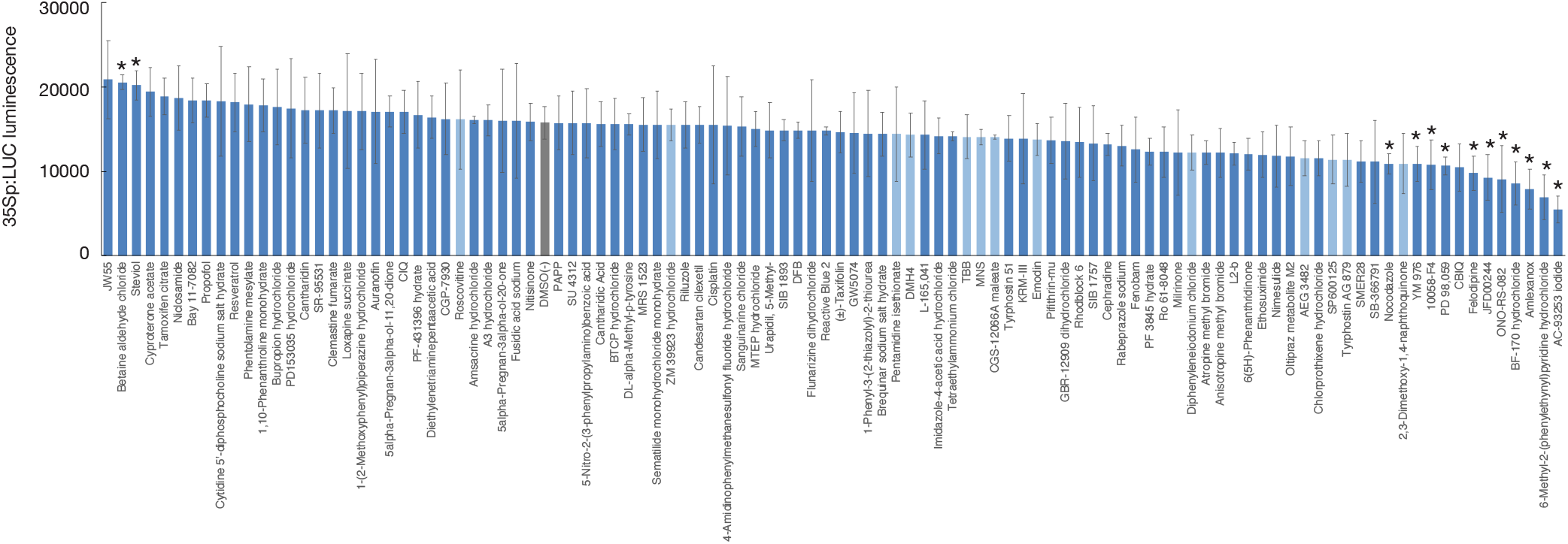
Inhibition of luciferase luminescence by LOPAC chemicals. Luciferase luminescence in *35Sp:LUC* seedlings, 6 h after transfer to media with 25 µM chemical (means ± SD, n = 3).

**Figure S2.**
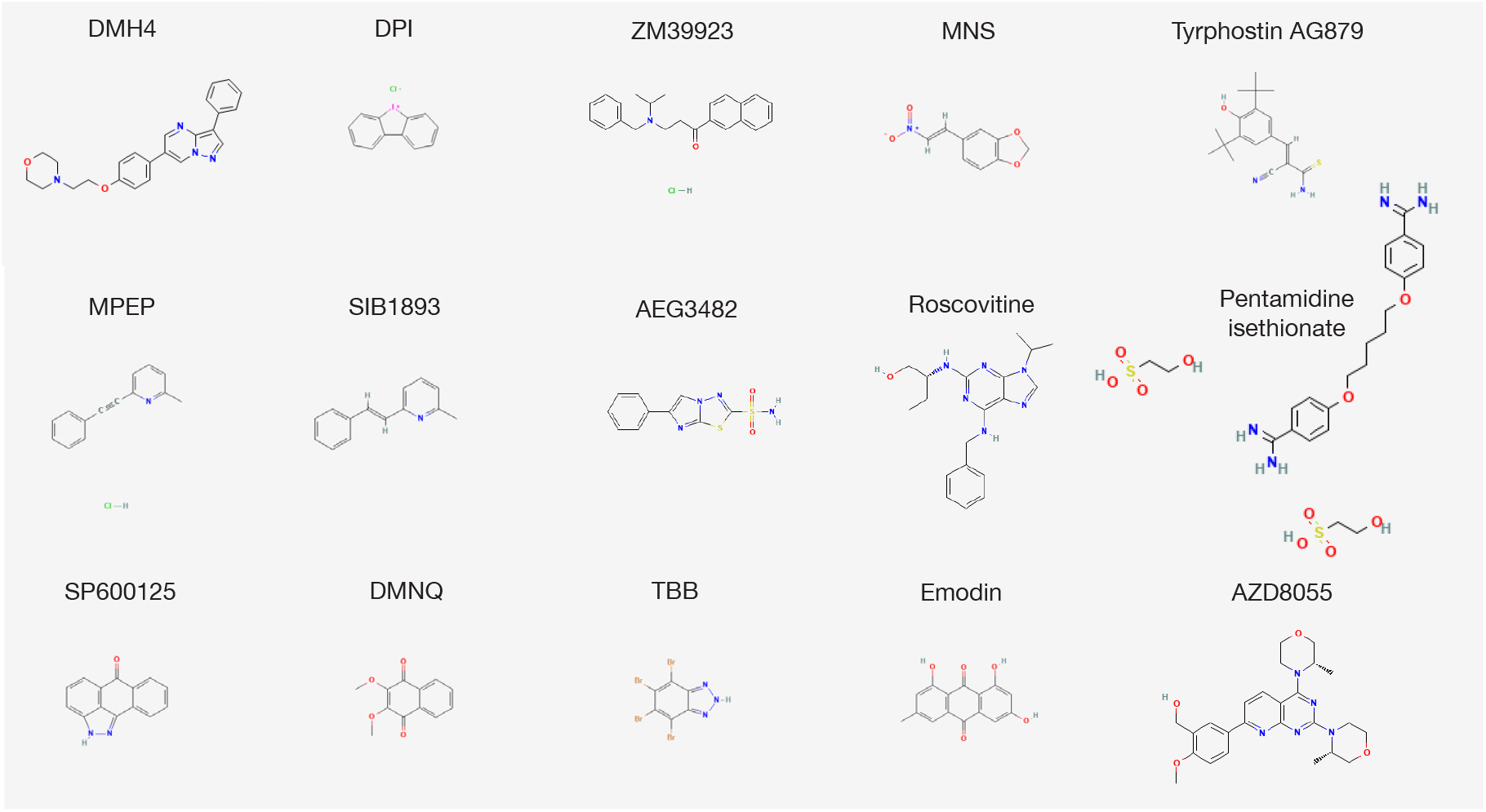
Structures of 15 chemicals chosen for validation. Chemical structures were obtained from Pubchem (https://pubchem.ncbi.nlm.nih.gov) and scaled to equivalent size.

**Figure S3.**
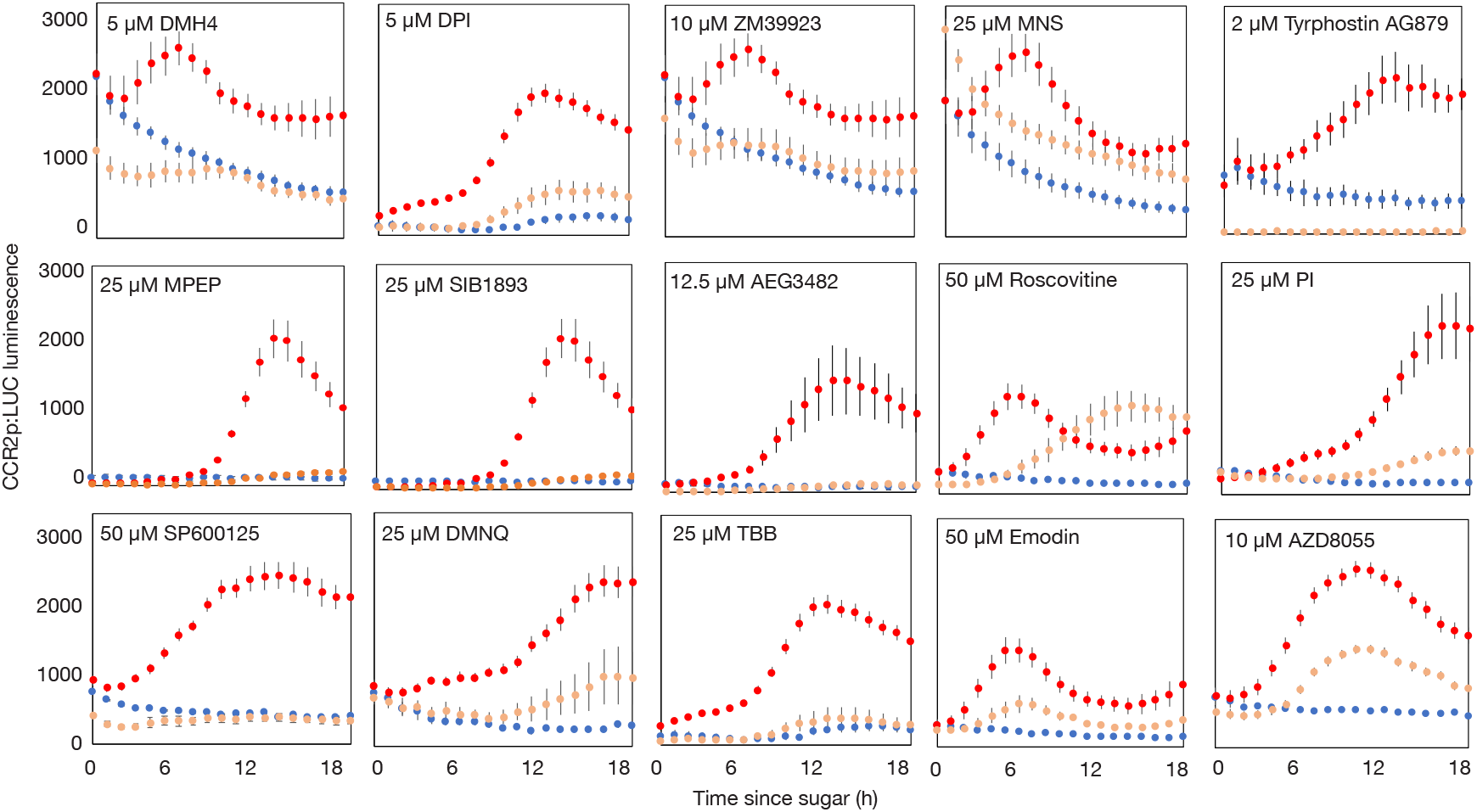
Inhibition of transcriptional response to sucrose. Luciferase luminescence in dark-adapted *CCR2p:LUC* seedlings following treatment with 30 mM mannitol (blue), 30 mM sucrose (red) or 30 mM sucrose and the indicated concentration of chemical (pink) (means ± SEM, n = 6-12).

**Figure S4.**
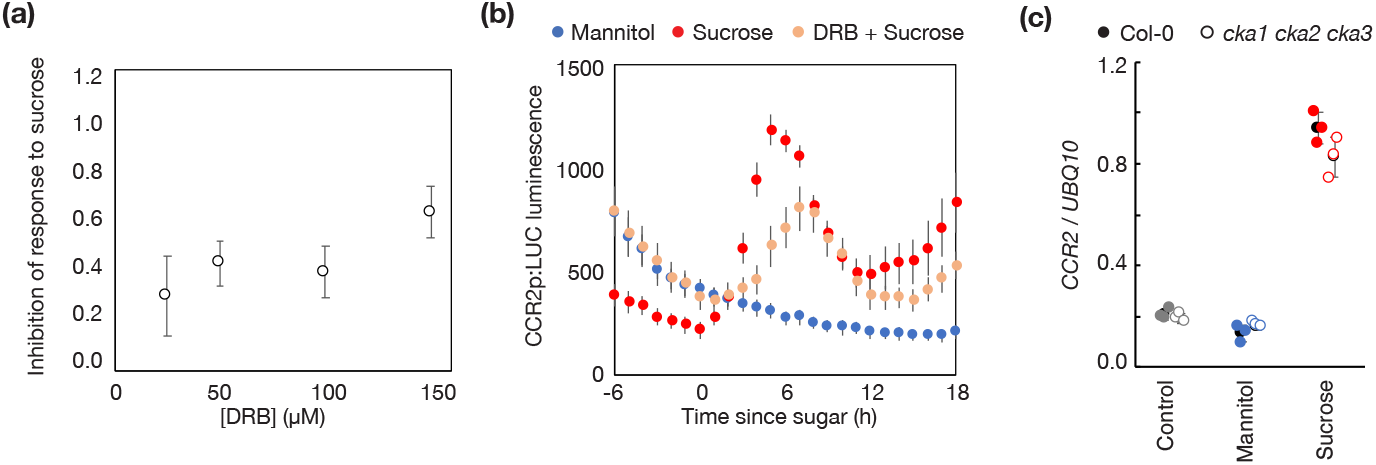
CK2 is not required for *CCR2* response to sucrose. (a) Inhibition of peak luciferase luminescence in dark-adapted *CCR2p:LUC* seedlings after treatment with 30 mM sucrose in the presence of the indicated concentration of DRB (means ± SD, n = 8). (b) Luciferase luminescence in dark-adapted *CCR2p:LUC* seedlings after treatment with 30 mM sucrose in the presence of 150 µM DRB (means ± SD, n = 8). (c) *CCR2* transcript level, relative to *UBQ10*, in dark-adapted wild-type (Col-0) and *cka1-1 cka2-1 cka3-1* seedlings before (Control) or 12 h after treatment with 30 mM mannitol or 30 mM sucrose (means ± SD, n = 3).

**Figure S5.**
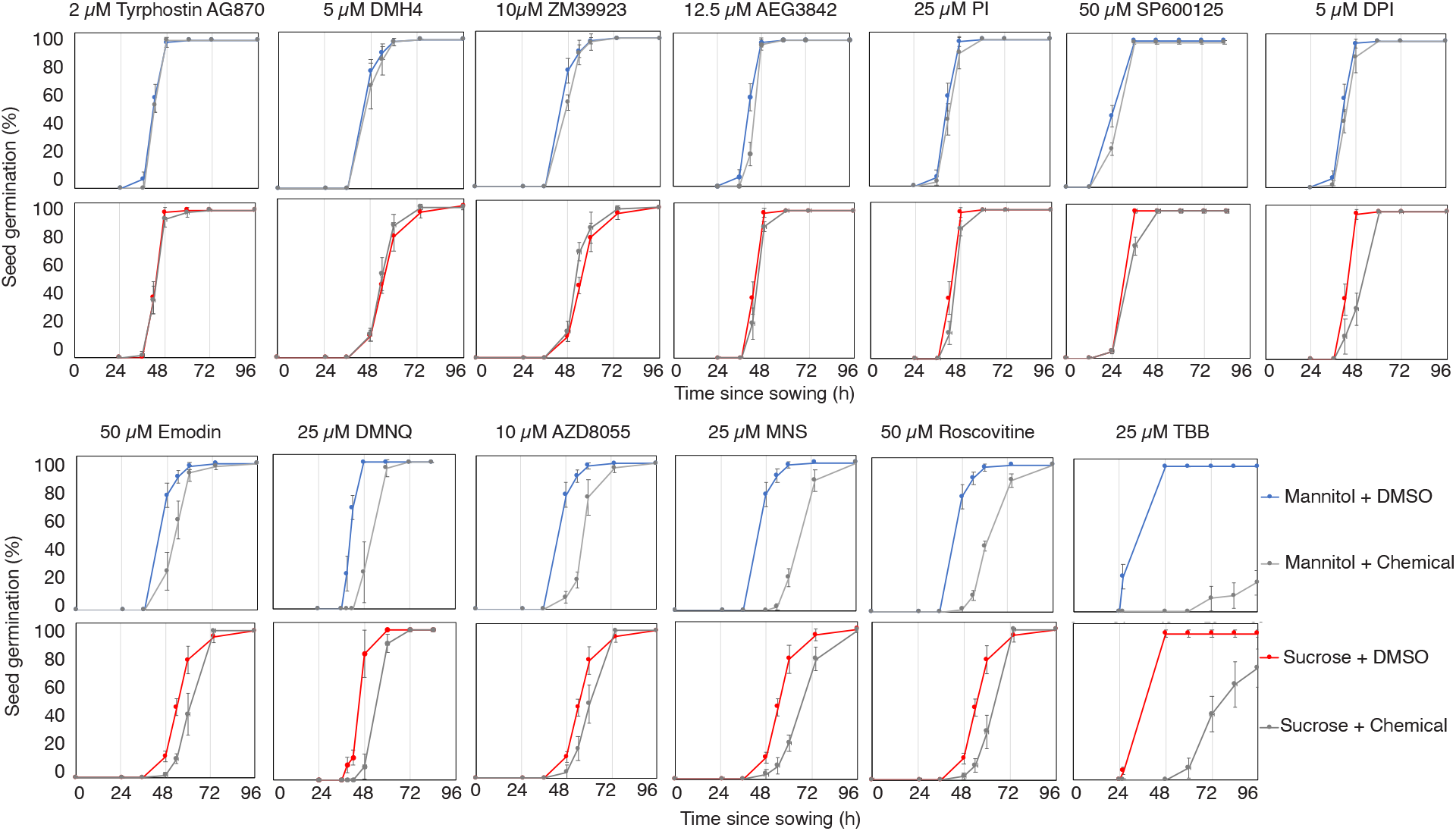
Effect of chemicals on germination of dormant seeds. Imbibed Col-0 seeds were sown on ½ MS containing 30 mM mannitol or 30 mM sucrose containing DMSO or indicated concentration of chemical (means ± SD of four replicate each of 20-25 seeds).

**Figure S6.**
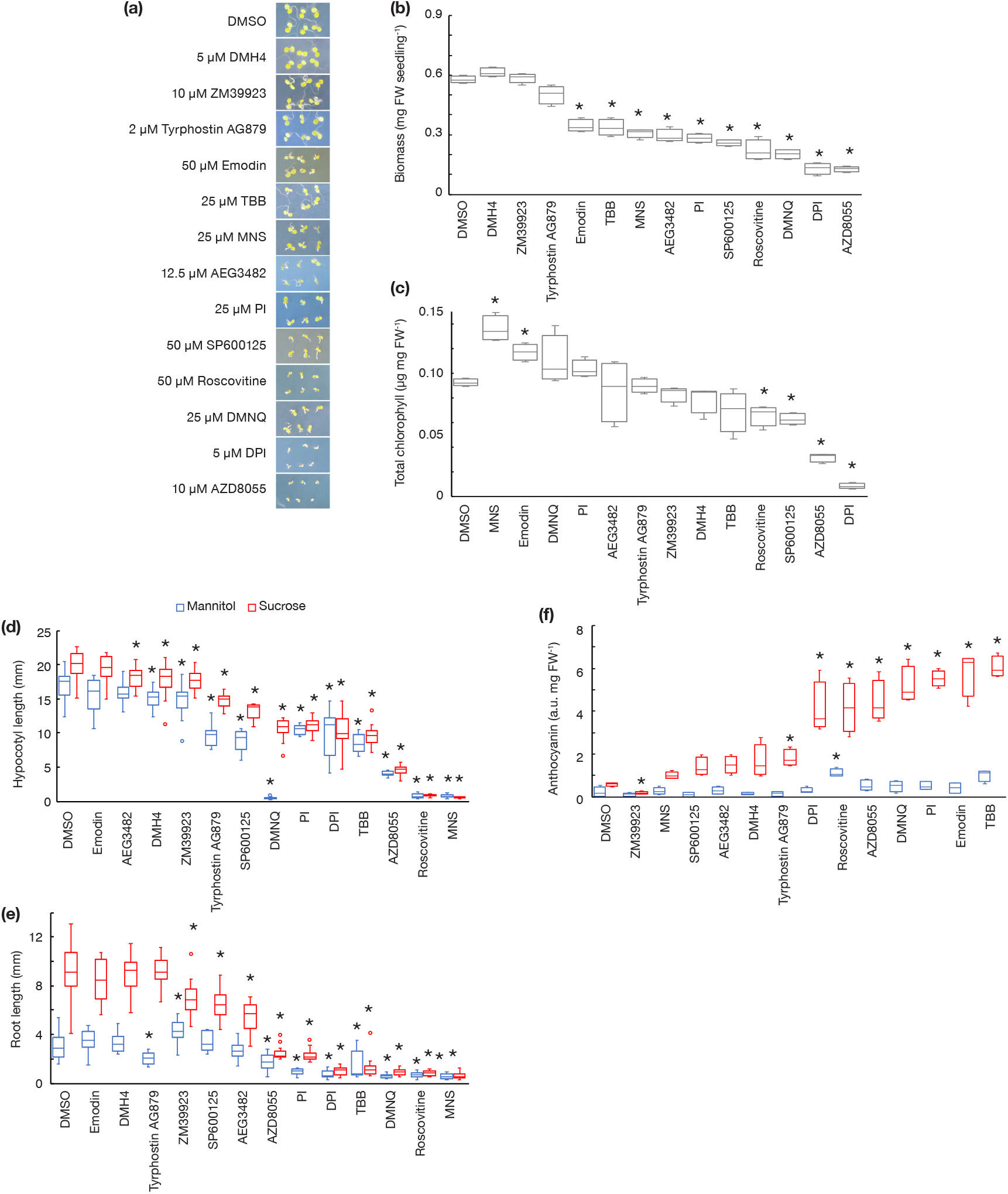
Effects of chemicals on growth and pigments. (a) Images, (b) fresh weight and (c) total chlorophyll of 7 d old seedlings grown on ½ MS containing DMSO or minimum effective concentration of each chemical (n=4). (d) Hypocotyl length and (e) root length of 7 d old seedlings grown in the dark for 5 d on 30 mM mannitol (blue) or sucrose (red) containing DMSO or chemical (n=10). (f) Anthocyanin content of 9 d old seedlings grown for 2 d on 90 mM mannitol (blue) or sucrose (red) with DMSO or chemical (n=4). * Different from control (DMSO) by Bonferroni-corrected *t*-test, p < 0.05.

**Figure S7.**
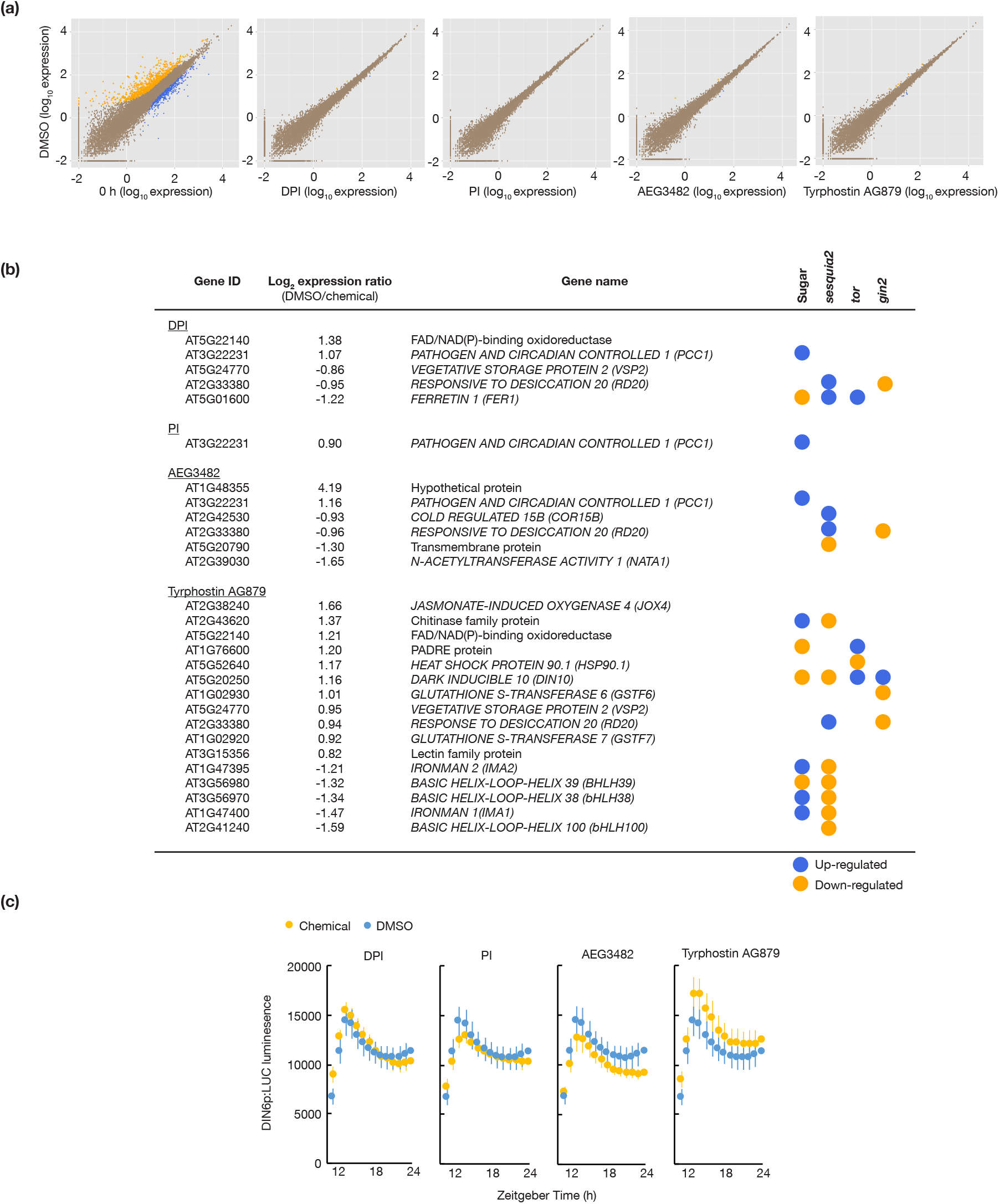
Effect of selected chemicals on transcriptome at dawn. (a) Comparison of gene expression in untreated seedlings at ZT0 (0 h) or after 2 h treatment with DMSO, DPI, PI, AEG3482 or Tyrphostin AG879. (b) List of differentially expressed genes between DMSO or chemical-treated seedlings. Genes previously identified as differentially expressed in dark-adapted seedlings treated with sucrose (Sugar; Román et al 2021) or in mutants of SnRK1 (*sesqui;* Peixoto et al 2021), TOR (*tor*; Xiong et al 2013) or HXK1 (*gin2;* Ganpudi et al 2019) are indicated by yellow (down-regulated) or blue (up-regulated). (c) Luciferase luminescence in *DIN6p:LUC* seedlings treated with DMSO or chemical before dusk (ZT11) (means ± SEM, n = 6).

**Figure S8.**
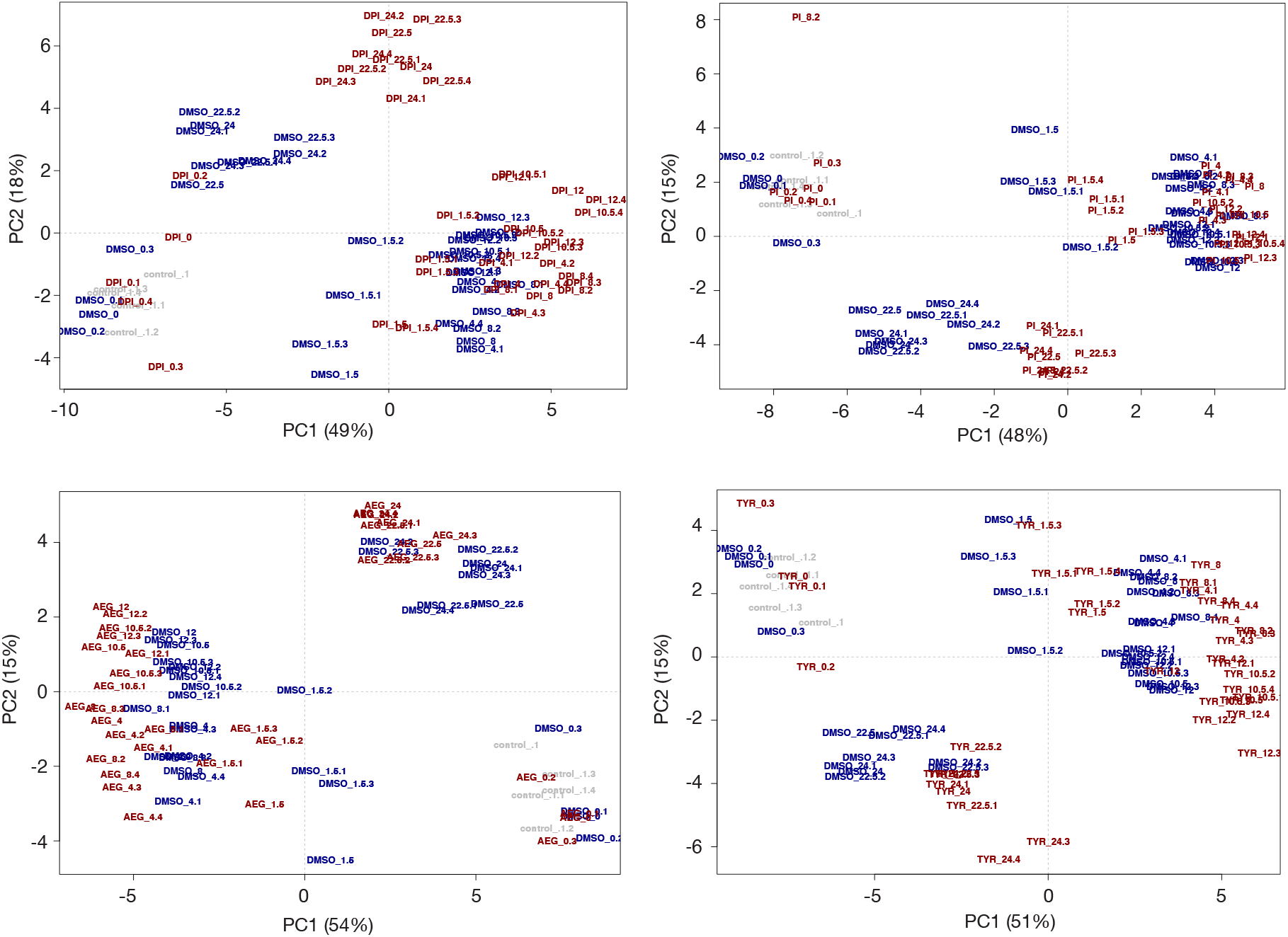
Principle component analysis of effects of chemicals on primary metabolites. Principle component 1 (PC1) versus PC2 plots for 63 metabolites measured in untreated seedlings at ZT0 or seedlings treated with DMSO, DPI, PI, AEG3482 (AEG) or Tyrphostin AG879 (TYR) and sampled at ZT0, ZT1.5, ZT4, ZT8, ZT10.5, ZT12, ZT22.5 and ZT24. Each point represents one of five biological replicates.

Table S1. Primers used in this study.

Table S2. LOPAC chemicals that modify *CCR2p:LUC* reporter response to sucrose, SSMD ±1.

Table S3. DEGs identified by RNA-seq in untreated seedlings or seedlings treated with DMSO, DPI, PI, AEG3482 or Tyrphostin AG879.

Table S4. Metabolite profiling in untreated seedlings or seedlings treated with DMSO, DPI, PI, AEG3482 or Tyrphostin AG879.

